# Mechanotransduction coordinates extracellular matrix protein homeostasis promoting longevity in *C. elegans*

**DOI:** 10.1101/2022.08.30.505802

**Authors:** Alina C. Teuscher, Cyril Statzer, Anita Goyala, Seraina A. Domenig, Ingmar Schoen, Max Hess, Alexander M. Hofer, Andrea Fossati, Viola Vogel, Orcun Goksel, Ruedi Aebersold, Collin Y. Ewald

## Abstract

Although it is postulated that dysfunctional extracellular matrices (ECM) drive aging and disease, how ECM integrity assures longevity is unknown. Here, using proteomics and *in-vivo* monitoring of fluorescently tagged ECM proteins, we systematically examined the ECM composition during *Caenorhabditis elegans* aging revealing three distinct collagen dynamics. We show that age-dependent stiffening of inert collagen was slowed by longevity interventions through prolonged replenishing of collagens. In genetic and automated lifespan screens for the regulators that drive this remodeling, we identify hemidesmosome-containing structures that span from the exoskeletal ECM through the hypodermis, basement membrane ECM, to the muscles, coupling mechanical forces to adjust ECM gene expression across tissues. The hemidesmosome tension-induced adaptation is mediated via transcriptional co-activator YAP. Our data reveal a novel mechanism of mechano-coupling and synchronizing of two functionally distinct and spatially distant ECMs that is indispensable for longevity. Thus, besides signaling molecules, mechanotransduction-coordinated ECM remodeling systemically promotes healthy aging.

**Graphical Abstract:** 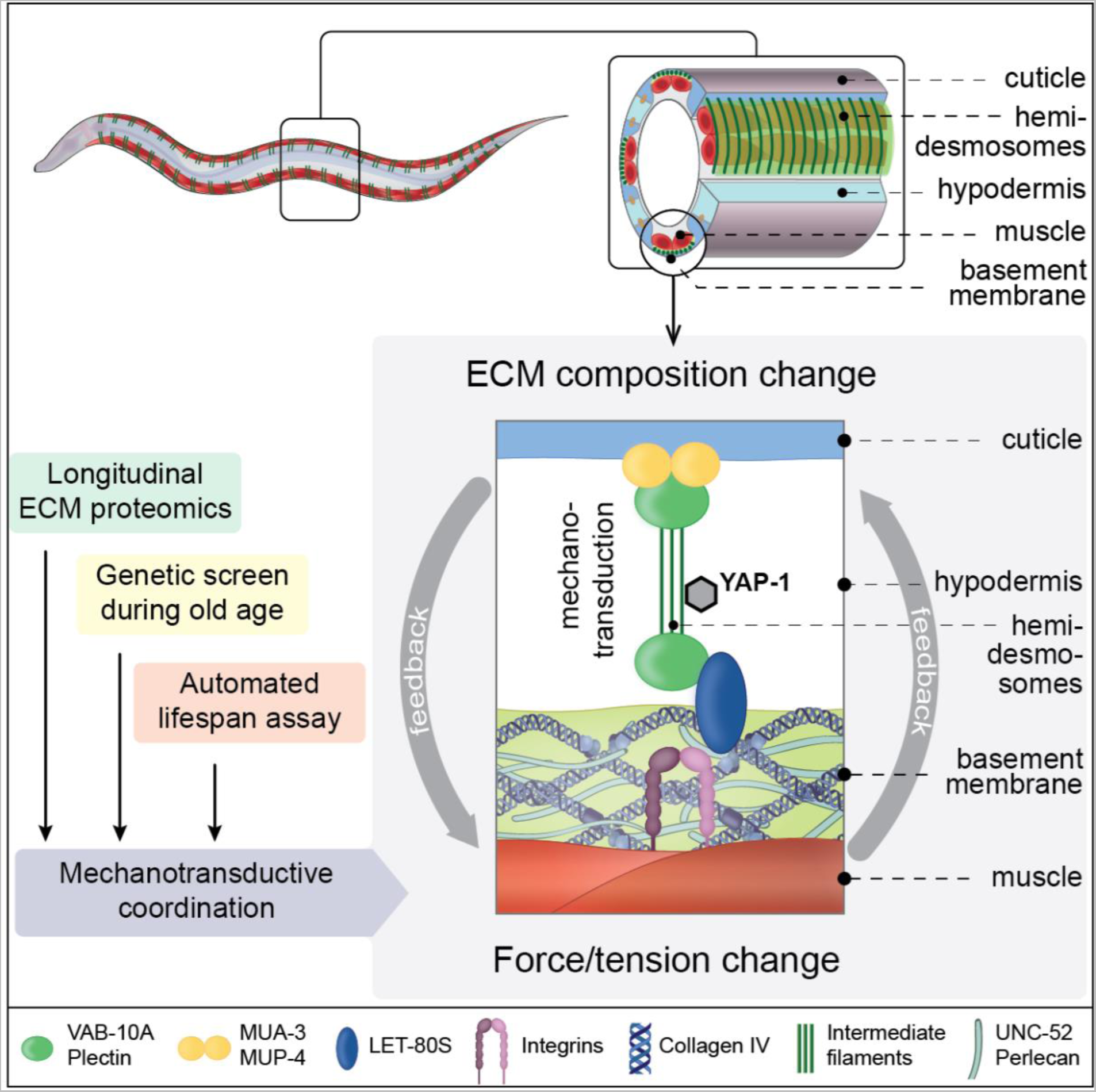

**Highlights:**

- Proteomics, genetics screen, and automated lifespan assays of >55’000 animals all point to hemidesmosome-containing structures for the mechano-regulation of ECM homeostasis and longevity
- Coupling of biomechanical properties of two ECMs with underlying cellular signaling
- Transcriptional co-activator YAP-1 is required for longevity and pressure-induced collagen homeostasis

## Introduction

Cells and organs are surrounded by and anchored to extracellular matrices (ECMs) (Bonnans et al., 2014; Frantz et al., 2010; Hynes, 2009). The ECM is a network composed of large multidomain proteins linked together to form stable structures essential for tissue geometry and integrity (Hynes, 2009). About 300 proteins, such as collagens, glycoproteins, and proteoglycans, form the actual matrix, the so-called core-matrisome (Naba et al., 2016). These proteins are produced, secreted, and incorporated to form the matrix (Lu et al., 2011). Each cell niche has the capability to produce its own ECM (Ewald, 2020; Frantz et al., 2010; McKee et al., 2019; Sacher et al., 2021). Thereby, each tissue is surrounded by its own unique ECM that entails unique physical properties (Humphrey et al., 2014).

The function of the ECM is to support and protect organs and tissues and facilitate inter-tissue communication (Hynes, 2009). To execute these functions, the ECM has recently been appreciated to be a highly dynamic structure while providing mechanical support (Vogel, 2018); it stores and presents growth factors and is enzymatically remodeled to assure cellular homeostasis (Bonnans et al., 2014; Ewald, 2020; Frantz et al., 2010; Hynes, 2009). These remodeling and signaling functions are performed and controlled by about 700 associated-matrisome proteins (Naba et al., 2016). The ∼1000 core- and associated matrisome genes comprise about 4% of the genome and are linked to more than ten thousand phenotypes encompassing about 10% of the entire phenome, including the developmental, structural, immune system, stress resilience, and age-related phenotypes in humans, mice, zebrafish, *Drosophila*, and *C. elegans* (Statzer and Ewald, 2020). Furthermore, the core matrisome proteins collagen type I, II, V, and glycoprotein fibrillin have each been associated with more than 150 distinct phenotypes, revealing the profound importance of a functional ECM (Statzer and Ewald, 2020).

Not surprisingly, changes in ECM composition are involved in many diseases, such as cancer, fibrosis, atherosclerosis, and neurodegenerative disorders (Ewald, 2020; Taha and Naba, 2019). For instance, based on their ECM composition cancer types can be identified, adverse patient outcomes can be predicted, as well as circulating tumor cells can be identified based on their ECM gene expression (LIM et al., 2017; Naba et al., 2014; Ting et al., 2014; Yuzhalin et al., 2018). Furthermore, deregulation of matrisome genes is a crucial step during the transition and reprogramming from normal healthy cells into tumor cells and for metastasis (Mitra et al., 2019). Thus, ECM composition does not only reflect cell identity but also their phenotypic state, health, or disease status. To conceptualize this, we named this phenomenon the ‘matreotype’. A matreotype is a ‘snapshot’ of the ECM composition associated with or caused by a phenotype or physiological states, such as health, disease, or aging (Ewald, 2020). Using RNA sequencing data, we have defined the youthful matreotype of humans, probed changes in gene expression upon drug treatment, and thereby predicted and validated several novel longevity drugs (Statzer et al., 2021a). This illustrates that the matreotype has broad implications for biomedical research. However, the underlying mechanism(s) of how changes in matreotype impact physiology are unknown.

Although RNA sequencing and proteomics can provide an indication of potential changes in ECM composition *ex vivo*, the functional consequences on cellular integrity and homeostasis *in vivo* during aging are largely unexplored. RNA levels rarely tightly correlate with protein levels (Liu et al., 2016), and the protein/mRNA abundance ratio is further complicated for proteins that form multicomplex structures, such as the ECM, where proteins are post-translationally processed, modified, secreted, incorporated, and crosslinked to form the matrix. To profile matreotype changes, we turned to the multicellular model organism *C. elegans,* which has two main extracellular matrices: the basement membrane, a sheet-like structure surrounding organs, and the cuticle that forms the exoskeleton (Kramer, 2005; Page and Johnstone, 2007). Its short lifespan of 3 weeks and the transparency of *C. elegans* allowed us to monitor fluorescently tagged ECM proteins incorporated into matrices non-invasively *in vivo* during aging.

Specifically, we monitor ECM proteins during aging to identify changes in matreotypes (*i.e.,* matrix composition). Since ECMs are bound via adhesion receptors such as integrins to cells and their cytoskeleton we also defined the adhesome of *C. elegans* to complement the matrisome. We show that certain cuticular collagens are remodeled out of the ECM during aging which is associated with a decline in the gene expression of these collagens. Longevity interventions prolong collagen expression and we utilize this finding for a targeted RNAi screen. We identify hemidesmosome-containing structures that encompass or connect many components of the basement membrane, integrin receptors, and adhesome-cytoskeleton signaling regulators underlying prolonged collagen homeostasis associated with longevity. Yes1 Associated Transcriptional Regulator (YAP-1) transforms the physical forces from hemidesmosome-containing structures to prolong collagens gene expression (*i.e.*, mechanotransduction) and longevity. We demonstrate that an age-dependent uncoupling of mechanotransduction abolishes the feedback, thereby inhibiting the prolonged ECM protein homeostasis and longevity. Thus, we provide new mechanistic evidence that mechano-coupling or mechano-transduction is essential for promoting healthy aging.

## Results

### ECM composition during aging

Our first goal was to determine whether ECM composition changes and remodels during aging. ECM remodeling starts with proteases, which excise and degrade proteins from the ECM. Excised proteins are then replaced by *de novo* synthesized ECM proteins that are secreted and incorporated into the matrix with the help of proteases and cross-linking enzymes (Lu et al., 2011). To capture this process, we assessed matrisome and adhesome dynamics by fluorescent reporters and generated new ECM-enriched proteomics data along an aging timeline, and combined them with previously published omics data on five different levels: (1) gene expression via RNA sequencing, (2) timing and localization of expression via promoter reporters *in vivo*, (3) matrisome protein levels via quantitative proteomics, (4) *de novo* synthesis of matrisome proteins based on SILAC-label-chase proteomics data, and (5) monitoring of selected matrisome proteins tagged with fluorescent proteins incorporated into the ECM *in vivo* during aging (Figure 1 and Supplementary Figure 1).

**Figure 1.**
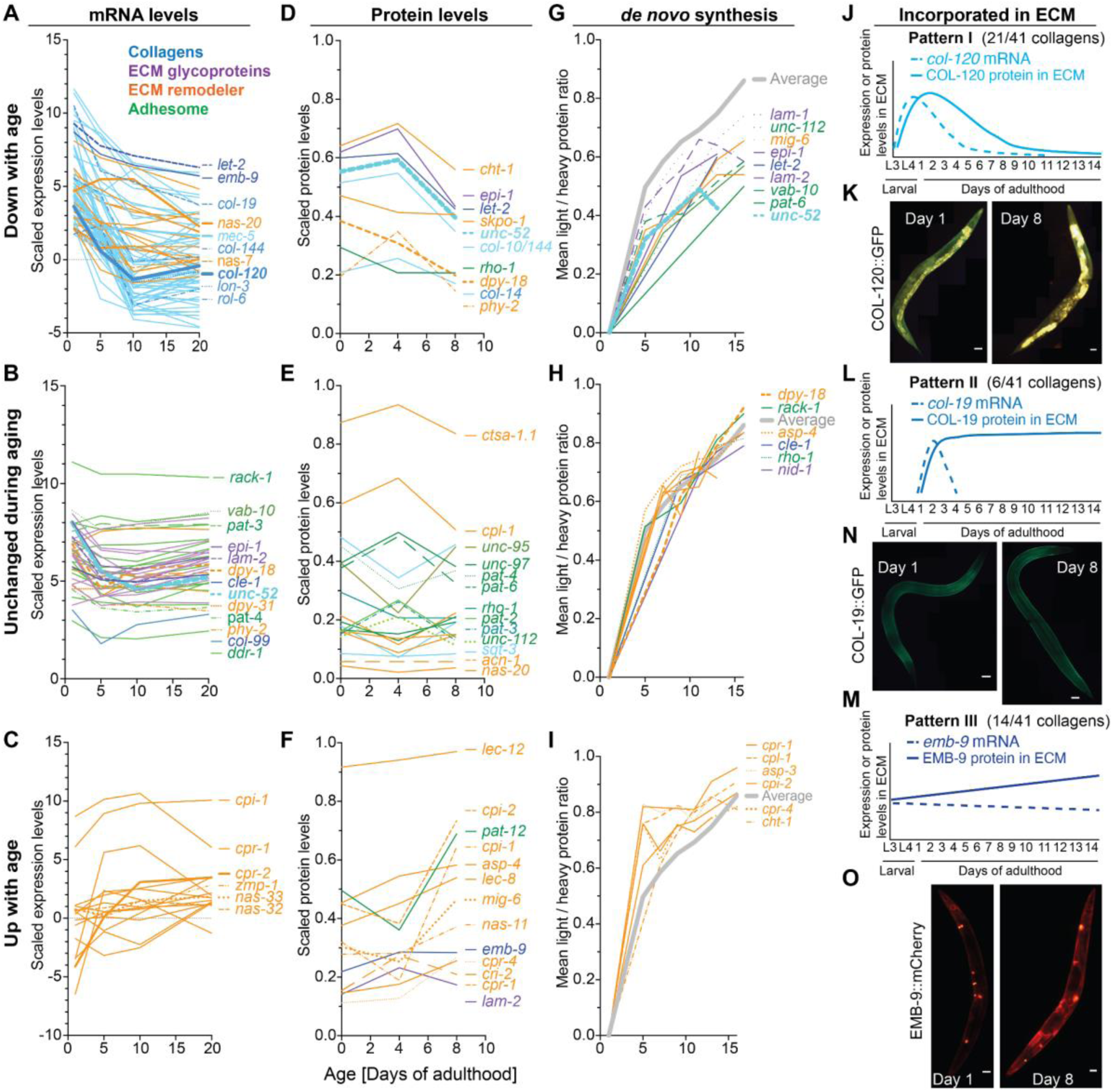
Matrisome dynamics during aging. (A-C) Aging time course of matrisome mRNA levels were classified as down- (A), unchanged (B), or up-regulated (C) during aging based on the agreement between the Pearson and Spearman correlation coefficients when applied to the individual samples. (Source data: GSE46051, Supplementary Table 2). (D-F) ECM-enriched proteomics aging time course matrisome protein levels declining (A), unchanged (B), or increasing (C) during aging taking into account the aging correlation of the individual samples (Data and statistics Supplementary Table 3). (G-I) Classification of the *de novo* protein synthesis rate of each protein into lower (G), unchanged (H), or elevated (I) during aging based on the difference between the synthesis rate of the individual protein to the overall mean protein production. (Source data: PMID: 25686393, Supplementary Table 3). (J-O) The aging time course of collagens that are incorporated into the ECM. (J, L, M) Model extrapolated from transgenic fluorophore tagged collagens shown as representative images (K, N, O). Details in Supplementary Figure 1, Supplementary Table 1. Scale bar = 50 µm.

Starting in early adulthood, during aging the majority of cuticular collagen (*col*) mRNA levels steeply declined, accompanied by a decline in some molting-associated ECM remodelers (protease/*nas*) (Figure 1A, Supplementary Figure 1I, 2, Supplementary Table 1, 2) (Budovskaya et al., 2008; Schmeisser et al., 2013). The mRNA levels of adhesome (integrin/*pat-3*), ECM glycoproteins (laminin/*lam-2/epi-1*), some conserved collagens (Type XVIII/*cle-1,* Type XXV/*col-99*) accompanied by some pro-collagen processing and stabilization enzymes (procollagen C-peptidase/*dpy-31*, prolyl 4- hydroxylase/*dpy-18/phy-2*) were unchanged and continuously expressed during aging (Figure 1B, Supplementary Figure 1I, Supplementary Table 1, 2). By contrast, the only category of matrisome genes that increased in expression during aging were proteases and protease inhibitors that remodel the ECM, such as MMP/*zmp*, astacin metalloprotease/*nas*, cathepsin/*cpr*, and protease inhibitor/cystatin/*cpi* (Figure 1C, Supplementary Figure 1I, Supplementary Table 1, 2).

**Figure 2.**
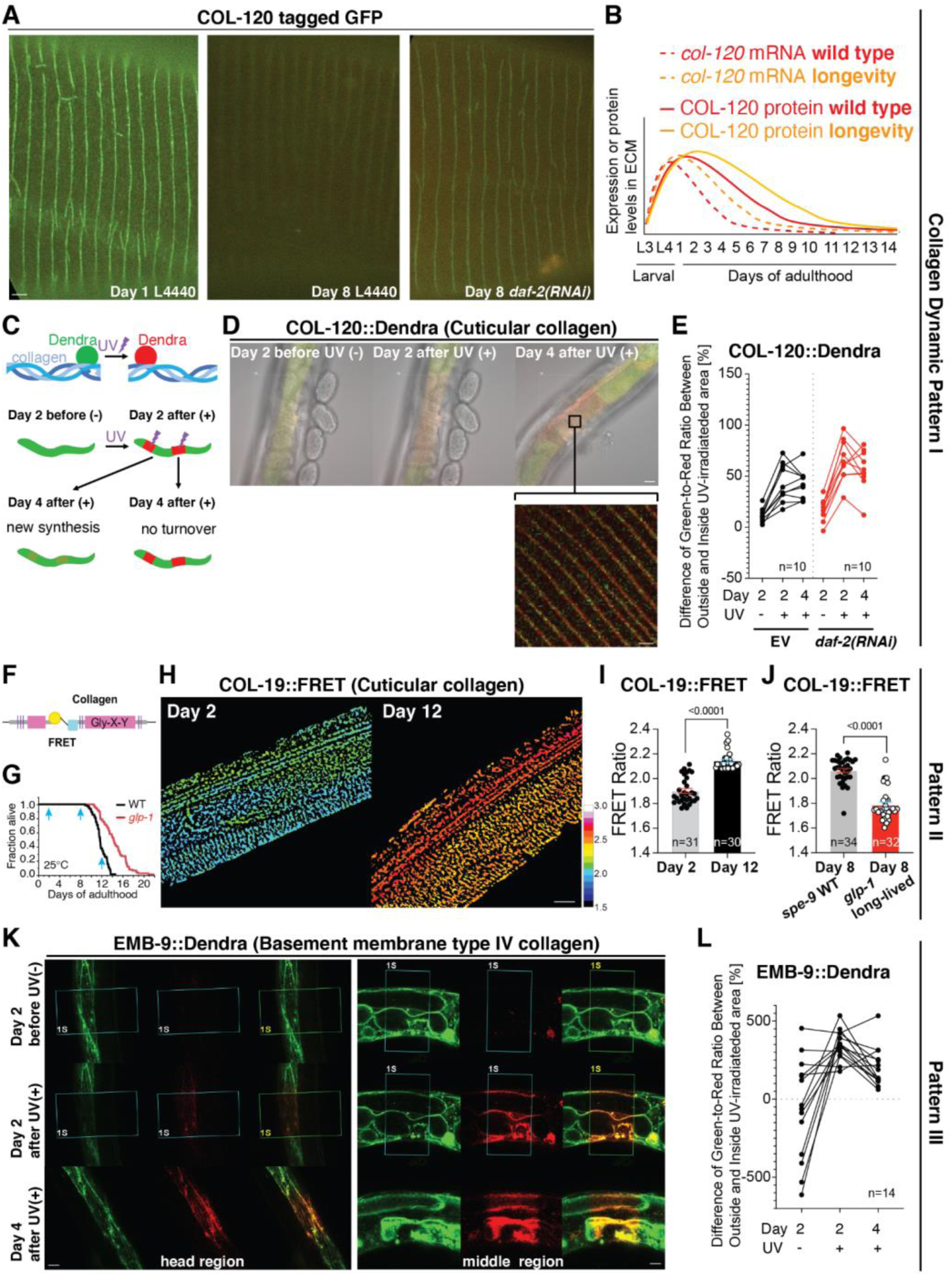
Three patterns of collagen dynamics during longevity quantified *in vivo*. (A-E) COL-120 dynamics as a representative for the pattern I collagen. (A) Collagen COL-120 tagged with GFP (LSD1000) in the cuticular ECM furrow vanished from day 1 to day 8 on control empty vector RNAi (L4440) but was still visible at day 8 of adulthood when *daf-2(RNAi)* started at L4. Scale bar = 1 µm. (B) Model of COL-120 mRNA, protein, and incorporated ECM reporter dynamics. © Experimental workflow and the expected outcome of photoswitching COL-120 tagged dendra2 *in vivo*. (D) Representative images of photo-switched areas of LSD1061 COL-120::Dendra Scale bars = 10 µm for overlay and 1 µm for inset. (E) Quantification of COL-120 turnover of the mid-body area. Each dot represents 1 animal (n=10). See Supplementary Table 5 for statistical details and raw data. (F-J) COL-19 is shown as a representative of the pattern II collagen. (F) Schematic representation of COL-19::FRET transgene. (G) Lifespan of wild-type temperature-sensitive sterile background WT (LSD2052 *spe- 9(hc88)*; COL-19::FRET) vs long-lived (LSD2053 *glp-1(e2141)*; COL-19::FRET) at 25°C. Light blue arrows indicate sampling days for FRET imaging. (H) Representative LSD2052 *spe-9(hc88)*; COL-19::FRET images in the 3D chamber on days 2 and 12 of adulthood at 25°C. The increasing FRET ratio is color-coded from dark blue to red. Scale bar = 10 µm. (I) FRET ratio increased from day 2 to 12 during aging of LSD2052 *spe-9(hc88)*; COL- 19::FRET at 25°C. See Supplementary Table 8 for statistical details and raw data. (J) Long-lived LSD2053 *glp-1(e2141)*; COL-19::FRET showed lower FRET ratio compared to WT (LSD2052 *spe-9(hc88)*; COL-19::FRET) at day 8 of adulthood at 25°C. See Supplementary Table 6 for statistical details and raw data. (K-L) Type IV collagen EMB-9 is shown as a representative of the pattern III collagen. (K) Representative images of photo-switched areas of NK860 EMB-9::Dendra. Scale bar = 10 µm. (L) Quantification of EMB-9::Dendra showed that newly synthesized EMB-9 is laid on top of the old matrix-incorporated EMB-9, which was not turnover. Each dot represents 1 animal (n=14). See Supplementary Table 8 for statistical details and raw data.

To determine the changes in ECM protein composition (*i.e.,* matreotype) during aging, we performed ECM-enriched proteomics on young (day 0 = L4), mature (day 4), and post-reproductive (day 8) *C. elegans.* On day 8 deaths from senescence were not observed. Mirroring the mRNA expression, the protein levels of several cuticular collagens declined during aging, adhesome protein levels were unchanged and proteases and protease inhibitors increased during aging (Figure 1D-F, Supplementary Table 3). By contrast, some ECM proteins increased with aging, although their mRNA levels declined (*e.g.,* collagen type IV/*emb-9*; Figure 1A, 1F). The protein levels of other matrisome genes that showed continuous mRNA expression declined during aging (*e.g.,* laminin/*epi-1*, perlecan/*unc-52*, prolyl 4-hydroxylase/*dpy-18/phy-2*; Figure 1B, 1D). This was consistent with lower *de novo* synthesis of these ECM proteins during aging (Figure 1G, 1H, Supplementary Table 3) (Vukoti et al., 2015). Consistent with the increased mRNA and protein levels, proteases and protease inhibitors maintained a higher *de novo* synthesis rate during aging (Figure 1I, Supplementary Table 3).

To assess the actual levels of proteins that are incorporated into the ECM, we followed fluorescent protein-tagged ECM proteins *in vivo*. The challenge was to place the tag in the protein sequence so it did not interfere with its function and incorporation into the ECM. Furthermore, the extremely low fluorescent signal of ECM tagged proteins is masked by the accumulation of autofluorescent waste products (*e.g.,* lipofuscin) during aging. In addition, isolated cuticles also start to auto-fluoresce during aging presumably due to accumulated glycation adducts on collagens (Davis et al., 1982). To overcome these challenges, we built a triple-band filter set to distinguish autofluorescence from ECM tagged GFP signals (Teuscher and Ewald, 2018). Based on our omics analysis, we assessed 21 tagged ECM proteins across the core-matrisome and adhesome categories during development and aging (Supplementary Figure 1J, Supplementary Table 1). We observed an age-dependent increase in abundance in basement membrane proteins, such as laminin/*lam,* perlecan/*unc-52*, collagen type IV/*emb-9*, and *myotactin-fibronectin- repeats/let-805* (Supplementary Figure 1J, Supplementary Table 1).

Collagens make up the majority of proteins in the ECM (Frantz et al., 2010). Out of the 181 *C. elegans* collagens (Teuscher et al., 2019a), we were able to assess the quantitative abundance data for 41 collagens proteins and mRNAs during aging (Supplementary Table 1). For these collagens, we observed three distinct dynamic patterns (I-III) during aging (Figure 1J-O, Supplementary Table 1). The pattern I consists of 21/41 detected collagens for which the mRNA, protein levels, and abundance in the ECM steeply declined during aging (*e.g, col-120*; Figure 1J-K, Supplementary Figure 1I- J, Supplementary Table 1-3). Pattern II consists of 6/41 detected collagens for which the mRNA steeply declined in early adulthood but the protein levels and/or abundance in the ECM stayed unchanged or increased during aging (*e.g, col-19*; Figure 1L-N, Supplementary Figure 1I-J, Supplementary Table 1-3). Pattern III consists of 14/41 detected collagens for which the mRNA remained unchanged or mildly declined but the protein levels and/or abundance in the ECM stayed increased during aging (*e.g, emb-9*; Figure 1M-O, Supplementary Figure 1I-J, Supplementary Table 1-3).

Taken together, we mapped the dynamic ECM composition (*i.e.,* matreotype) during aging. Our data indicate that some ECM components are once synthesized, incorporated, and stay lifelong in the ECM, whereas other components are excised from the ECM during aging, and yet other ECM components are continuously added to the ECM.

### Longevity interventions prolong collagen pattern I expression during aging

Next, we asked which of these three dynamic collagen patterns are altered *in vivo* upon longevity interventions. To slow aging, we used *daf-2(RNAi)* to reduce Insulin/IGF-1 receptor signaling. For pattern I, using promoter-driven transgenic animals, we observed that *daf-2(RNAi)* prolonged the expression of *col-120* mRNA during aging (Supplementary Figure 3A, 3B, Supplementary Table 4). While COL-120 protein tagged with GFP gradually disappeared from the cuticular ECM during aging, slowing aging by rIIS showed COL-120 in the ECM for a prolonged time (Figure 2A, 2B, Supplementary Figure 3C, 3D, Supplementary Table 4). Similar dynamics were observed with other collagens from the pattern I (Supplementary Figure 3E-J, Supplementary Table 4).

**Figure 3.**
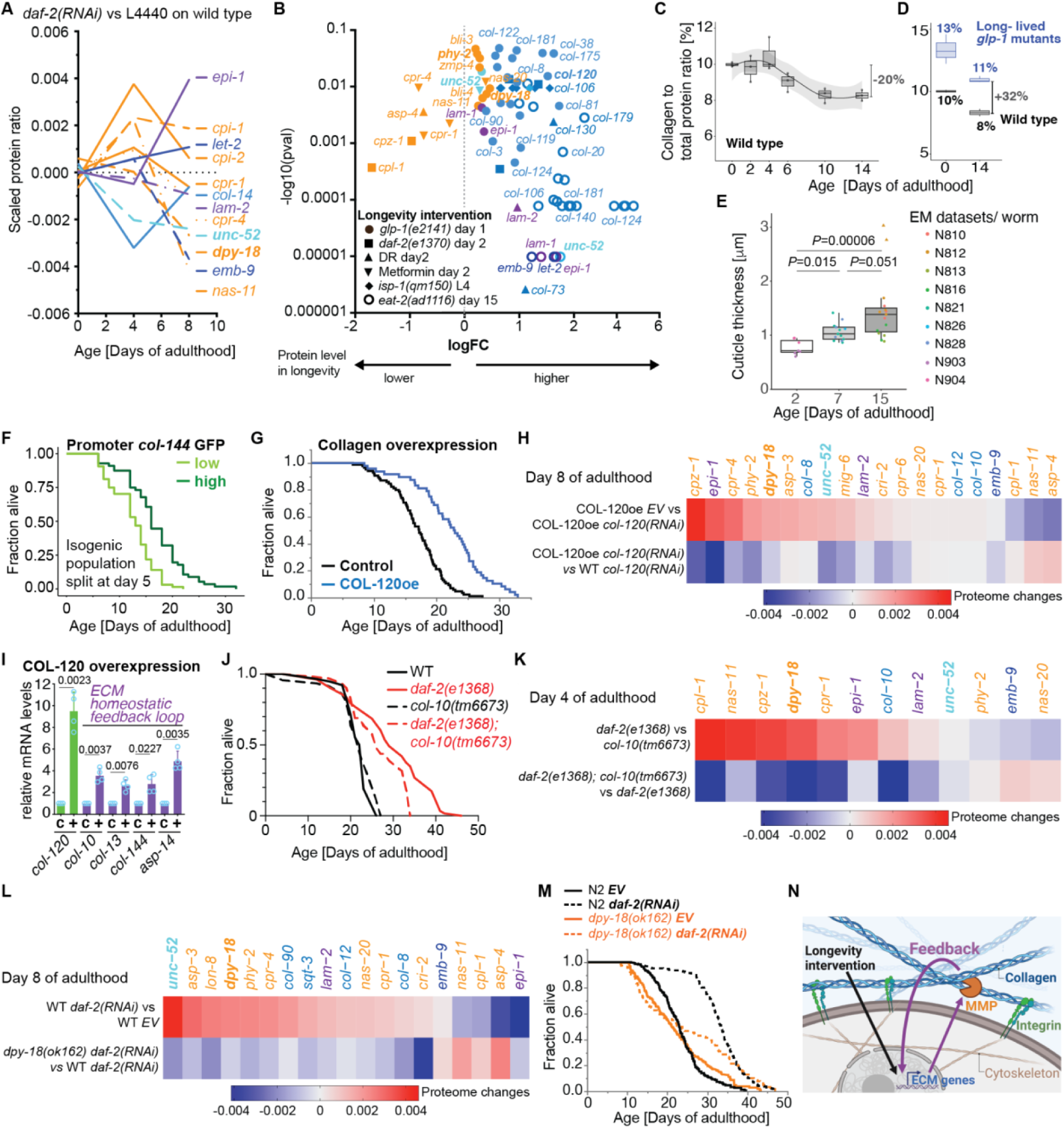
Feedback loop of ECM homeostasis implicated in longevity. (A) Time course of counterbalancing age-related changes in matrisome protein levels by *daf-2(RNAi)*-longevity intervention. Details in Supplementary Table 9. (B) Longevity interventions increased the normally age-related decrease of collagen levels and dampened the normally age-related elevation of extracellular proteases. Details in Supplementary Table 9. (C) The collagen over total protein content is displayed as a time course for a *spe-9 quasi wild type* population. (D) The collagen over total protein ratio is shown for *spe-9* and *glp-1* mutant populations at days 0 and 14 of adulthood. (E) Cuticle thickness increases with age based on electron microscopy (EM) images (Source: wormimage.org). Individual *C. elegans* are represented as dots (EM dataset). Triangles are the outliers. *P* values are One-way ANOVA post hoc Tuckey. See Supplementary Figure 5 and Supplementary Table 9 for details. (F) Isogenic population of *col-144* promoter GFP (LSD2002 *spe-9(hc88)*; P*col-144*::GFP) *C. elegans* were split at day 5 of adulthood into high and low expressing GFP individuals. (G) Collagen COL-120oe (LSD2017) overexpression increased lifespan compared to control (wild type with *rol-6(su1006)* co-injection marker LSD2013) on UV-inactivated bacteria. (H) Differences in protein abundance ratios are displayed for COL-120oe and wild-type populations undergoing control and *col-120* RNAi treatment. (I) Overexpression of COL-120oe (LSD2017) increased mRNA levels of other collagens and ECM proteases by qRT-PCR at day 1 of adulthood. N=4 independent biological samples in duplicates (each over 200 L4 worms). Mean ± SEM. *P* values relative to WT were determined by a one-sample t-test, two-tailed, with a hypothetical mean of 1. (J) Collagen *col-10(tm6673)* mutation partially suppressed *daf-2(e1368)* reduced insulin/IGF-1 receptor signaling longevity at 20°C. (K) Changes in protein abundance ratios are show for *daf-2(e1368)*, *col-10(tm6673)*, and *daf-2(e1368); col-10(tm6673)* animals. (L) The effect of *daf-2* RNAi on changes in protein abundance ratios is shown in wild type and *dpy-18(ok162)* populations. (M) Prolyl 4-hydroxylase *spy-18(ok162)* mutation partially suppressed reduced insulin/IGF-1 receptor signaling longevity upon adulthood-specific knockdown of *daf-2* at 20°C. (N) Model of the extracellular matrix homeostasis feedback loop. (F, G, J, M) For details, raw data, and statistics, see Supplementary Table 7.

To assess protein turnover, we tagged COL-120 with Dendra, a photoswitchable fluorophore (Dhondt et al., 2016; Ihara et al., 2011). Because COL-120 starts to disappear from the ECM in early adulthood, we irreversibly photoconverted COL-120::Dendra from green to red fluorescence on day 2 of adulthood, let the animals age for two more days, and then assessed green versus red fluorescent COL-120::Dendra in the cuticle (Figure 2C). If during these two days no new COL-120 would be synthesized, then the photoconverted area would stay red. If all photoconverted COL-120 would be replaced (*i.e,* turned over), the photoconverted area would turn green. We found that the photoconverted areas mostly stayed red but new COL-120 collagens (in green) were added on top of the older COL-120 (in red; Figure 2D, Supplementary Table 5). Since COL-120 levels gradually decline in the ECM during this time period, we included the levels of the outside regions to subtract this general decline from quantification in the photoconverted area. We found that the old COL-120 disappeared faster from the ECM faster than the new COL-120 was added (Figure 2E, Supplementary Table 5). Regarding the dependency on longevity, slowing aging by *daf-2(RNAi)* enhanced and prolonged the addition of newly synthesized COL-120 onto the old COL-120 in the ECM (Figure 2E, Supplementary Table 5). Our representative example suggests that longevity interventions might counteract the gradual loss of pattern I collagens from the cuticle by simply adding newly synthesized collagens to the older collagens that are continuously excised out of the cuticle. This is consistent with the prolonged mRNA expression of these collagens and the higher collagen protein levels of longevity interventions during aging.

### Age-dependent loss of mechanical tension of stably intercalated pattern II collagens is rescued by longevity interventions

As COL-19 is a representative member of the pattern II collagens, we next assessed its turnover during aging. The *col-19* mRNA declined rapidly during early adulthood, but the GFP tagged COL-19 stayed incorporated during aging. Upon *daf-2(RNAi),* the cuticular COL-19 protein levels in the ECM compared to control remained unchanged during aging (Supplementary Figure 3I-J, Supplementary Table 4), suggesting that these collagens once synthesized and incorporated into the cuticle would stay lifelong in this ECM.

During aging, collagens that are not replaced accumulate advanced glycation end products (AGEs) leading to crosslinking of collagens and stiffening of ECM (Ewald, 2020). Isolated *C. elegans* cuticles become stiffer with age (Rahimi et al., 2022) and show a marked increase in fluorescent spectral peaks reminiscent of AGE (Davis et al., 1982). Based on this and our observation that COL-19 protein stayed incorporated in the ECM during aging, we hypothesized that COL-19 might become crosslinked, thereby altering mechanical properties. To test this, we used a Förster resonance energy transfer (FRET) sensor incorporated into COL-19 (Figure 2F) that has been previously used to read out mechanical stress and forces (Meng et al., 2011). As expected for a mechanosensor, the FRET transmission of this COL-19::FRET increased when the animals were compressed between two coverslips (Supplementary Figure 4A). To avoid external forces, we built a flow chamber for FRET measurements (Supplementary Figure 4B). We scored FRET ratios when animals were young (day 2 of adulthood), old but before death events occurred (day 8), and at very old age, when about 75% of the population had died (day 12; Figure 2G, Supplementary Figure 4C-F). We found a stark increase in FRET transmission during old age (day 12) compared to young (day 2) and this age-dependent increase in FRET transmission was lower in long-lived *glp-1* mutants at day 8 of adulthood compared to normal-lived wild-type controls (Figure 2H-J, Supplementary Table 4). This suggests that longevity interventions counteract age-dependent crosslinking of pattern II collagens.

To assess whether the increase in FRET transmission of this COL-19::FRET sensor corresponds to a reduction in its extensibility due to crosslinking, we fixed *C. elegans* with the crosslinking-agent formaldehyde which increased the FRET transmission compared to anesthetized animals (Supplementary Figure 4G-H). However, treating *C. elegans* with agents that either increase or decrease AGEs had minor effects on COL-19::FRET ratios and on lifespan (Supplementary Figure 4I-Q, Supplementary Table 7), arguing against collagen crosslinking as the sole driver for the age-dependent increase in FRET transmission.

An alternative reason for age-dependent changes in FRET transmission could be a change in tissue tension. Tissue tension is established and maintained by cells pulling on the ECM, either within the tissue itself or in neighboring tissues. To assess tissue tension, we used sodium chloride to remove the internal osmotic pressure leading to wrinkling of the cuticle at young (day 2) and old (day 8) age. We found that the age-dependent increase of COL-19::FRET transmission was nullified by loss of internal pressure in wild type and long-lived *glp-1* animals at day 8 of adulthood (Supplementary Figure 4R-U). However, young animals still had lower FRET ratios after salt treatment, suggesting that not all of it is due to tissue tension but some part might be due to collagen cross-linking. Our observation is consistent with a recent finding that under osmotic-shock-induced shrinkage of *C. elegans*, longevity interventions prevent the age-dependent increase of cuticular stiffness, which is nullified by knocking down pattern I collagen *col-120* (Rahimi et al., 2022), strengthening our model that longevity interventions promote ECM homeostasis to also counteract collagen crosslinking.

We conclude that age-related changes including collagen crosslinking occur on collagens that have been synthesized during youth and stay inert integrated into the ECM. Cuticle integrity declines during aging in part due to the loss of cells adhering to the ECM and progressive loss of tissue tension, which is slowed by longevity intervention.

### Longevity interventions slow the age-dependent accumulation of basement membrane collagens

As type IV collagen EMB-9 is a representative member of the pattern III collagens, we examined EMB-9 tagged with Dendra. As before, we photoconverted on day 2 of adulthood and two days later quantified the green (new) to the red (old) ratio of this basement membrane collagen. The newly synthesized EMB-9 collagens were added to the old collagens resulting in a thickening of the basement membrane independent of *daf- 2(RNAi)* longevity interventions (Figure 2K, 2I, Supplementary Figure 3K, 3L), Supplementary Table 4, 8). This observation is consistent with the thickening of human basement membranes (up to 100 fold) during aging (Halfter et al., 2015).

### Longevity interventions counteract age-dependent ECM compositional changes

To elicit what constitutes a youthful matreotype or ECM composition upon longevity interventions, we treated wild-type animals with *daf-2(RNAi)* to slow aging and compared the protein levels relative to control using proteomic data acquired at different time points of aging (Figure 3A, Supplementary Table 9). We searched the longitudinal abundance data for signatures that might reinstate a youthful matreotype. We found that protein levels of laminin/*epi-1*, collagen type IV/*let-2*, prolyl 4-hydroxylase/*dpy-18, and* perlecan/*unc-52,* which normally decline during aging, were increased in long-lived *daf- 2(RNAi)* animals (Figure 3A, Supplementary Table 9). Furthermore, several proteases that are normally being elevated during aging were reduced, particularly starting during mid-age in long-lived *daf-2(RNAi)* animals (Figure 3A, Supplementary Table 9). Consistent with our proteomics, across six different longevity interventions (dietary restriction, metformin, *glp-1, daf-2, isp-1, eat-2*) and data sets (Depuydt et al., 2013; Espada et al., 2020; Jung et al., 2021; Koyuncu et al., 2021; Pu et al., 2017), we observed an increase of a subset of cuticular collagens (*col-*) protein levels, collagen-stabilizing and remodeling enzymes (*dpy-18, phy-2, bli-, nas-, zmp-*) and a decrease of cathepsin (*cpl-, cpz-, cpr-*) protease levels (Figure 3B, Supplementary Table 9). We thus propose that longevity interventions mobilize compensatory adjustments counteracting age-related ECM changes, especially in older animals, presumably to maintain homeostasis of the ECM proteins.

Given that the largest observed changes occurred with enzymes that remodel collagens and with collagens, we quantified the overall collagen levels during aging. We found that one-fifth of the total collagen mass normalized to total protein mass is lost during aging and that longevity interventions started with more collagen mass which declined at a similar rate during aging (Figure 3C, 3D). This is in line with previous observations that longevity interventions have higher collagen levels during old age (Ewald et al., 2015). The cuticle is the fifth largest body part of *C. elegans,* making up about one hundred thousand µm^3^ or 1/6th of the total volume (Froehlich et al., 2021).

Because many cuticular collagens decline during aging, we reasoned that there might be a thinning of the cuticle occurring and thereby a loss of barrier protection, similar to the age-dependent loss of collagen and thinning of the human skin (Shuster et al., 1975). However, based on electron microscope pictures, the cuticle thickened in total by 0.197 µm (18%) during old age (day 7-15 of adulthood; Figure 3E, Supplementary Figure 5), consistent with previous observations (Herndon et al., 2002; Wolkow et al., 2017). It is unclear what underlies the thickening of the cuticle during aging, but given the massive decline in collagen levels, it might be other cuticular components such as the insoluble cuticulins, an accumulation of water, or a loosening of structural integrity as observed by EM (Essmann et al., 2020).

We conclude that the pattern I collagens are remodeled out of the ECM during aging, accounting for a gradual loss of collagen mass which is counterbalanced by longevity interventions.

### A feedback loop is sufficient and required for ECM homeostasis and longevity

Our observed time course of the collagen mass changes coincides with the growth rates in body size during adulthood, whereby after the final molt from L4 to adult, *C. elegans* continuously grows until day 6-8 of adulthood and then starts to shrink (Hulme et al., 2010; Shi et al., 2017; Statzer et al., 2022). *C. elegans* growth during early adulthood means an extension of the cuticular exoskeleton. We have previously shown that longevity interventions prolong this adult growth phase compared to wild type (Statzer et al., 2022). The longer this adulthood growth phase is, the longer-lived is an individual *C. elegans* (Hulme et al., 2010). To test whether this correlation is associated with prolonged production of the pattern I collagens, we used transgenic animals expressing GFP driven by the collagen *col-144* promoter, whose expression gradually declines during aging (Ewald, 2021; Statzer et al., 2021a), and split this isogenic population grown in the same environment into high expression and low expression of P*col-144*::GFP transgenic animals at day 5 of adulthood. Animals expressing higher levels of P*col-144*::GFP at day 5 of adulthood lived longer than their genetically identical siblings with lower levels of *col- 144*-driven GFP expressed (Figure 3F, Supplementary Table7), demonstrating that prolonged pattern I collagen expression is associated with longevity.

Previously we had shown that overexpression of pattern I collagen is sufficient to increase lifespan. Further, adulthood-specific knockdown of some pattern I collagens collapsed the higher collagen mass and blocked lifespan extension across conserved interventions (Ewald et al., 2015). However, the underlying mechanism remains unknown. Given that *C. elegans* die from age-associated infection of proliferating *E. coli*, its primary food source, we measured the lifespan of *C. elegans* overexpressing collagen COL-120 on heat-killed bacteria. We confirmed that overexpressing COL-120 extended *C. elegans* lifespan on dead bacteria (Figure 3G, Supplementary Table 7), excluding the idea that higher cuticular collagen levels would extend lifespan by improving barrier function. Selecting collagens from our omics dataset that were unaltered upon longevity interventions and overexpressing them was not sufficient to increase lifespan (Supplementary Figure 6, Supplementary Table 7) (Ewald et al., 2015), suggesting the unique properties of longevity-promoting collagens.

To identify downstream mechanisms mobilized by collagen overexpression, we used COL-120 overexpressing (COL-120OE) animals and treated them either with a control empty vector (EV) or *col-120(RNAi)* from L4 and performed proteomic analyses on day 8 of adulthood. We found abundance changes in proteins governing the cytoskeleton dynamics (*tbcb-1*/Tubulin-specific chaperone B, *pat-6*/parvin, *ifd- 2*/intermediate filament), as well as proteins involved in pathogen and oxidative stress response, and metabolism, respectively (Supplementary Table 9). Interestingly, we observed changes enhancing ECM composition (*i.e.,* matreotype) that were specific to COL-120 overexpression (Figure 3H, Supplementary Table 9). These include enhancement of cuticular collagens, enzymes that remodel cuticles and stabilize collagen, basement membrane components, as well as reduction of some age-dependent upregulated proteases (Figure 3H). We confirmed by qRT-PCR that overexpression of COL-120 leads to upregulation of transcripts coding for other collagens and ECM-remodeling enzymes (Figure 3I), suggesting a feedback loop between ECM composition and ECM production by cells. This suggests a model that the abundance of some key collagens is read out to adjust the abundance of other ECM components, an important feature for assembling a functional matrix.

To identify the extent of ECM remodeling and downstream pathways, we compared *daf-2(e1368)* with and without *col-10* at day 4 of adulthood. As loss of *col-10* blunts the longevity of *daf-2* (Figure 3J, Supplementary Table 7), we found that the enhancement of metabolism, detoxification, and stress defense depended on *col-10* but also on changes in the cytoskeleton (Supplementary Table 9). For the matrisome changes, the most significant enhancement of *daf-2*-induced longevity depending on *col- 10* was with prolyl 4-hydroxylase/*dpy-18,* an enzyme important for collagen stabilization (Figure 3K, Supplementary Table 9), reinforcing the idea of a feedback loop and demonstrating the importance to remodel the ECM for the cellular reprogramming upon longevity interventions.

To further test this idea, we compared wild type and a P4H *dpy-18(ok162)* loss-of-function mutant treated either with control or *daf-2(RNAi)* from L4 and performed ECM-enriched proteomics at day 8 of adulthood (Figure 3L, Supplementary Table 9). We found that the longevity-matreotype upon *daf-2(RNAi)* was reverted to the aging-matreotype of wild type in the *daf-2(RNAi)*-treated P4H *dpy-18(ok162)* mutants (Figure 3L). Consistent with these findings P4H function did not affect wild-type lifespan, but the enhancement of P4H during aging was important for longevity. The *daf-2(RNAi)-*mediated longevity was abolished in P4H *dpy-18(ok162)* mutants (Figure 3M, Supplementary Table 7), demonstrating that collagen stability is required for the dynamic change of ECM composition associated with healthy aging and for lifespan extension.

In summary, these results show that longevity interventions prolong the dynamic ECM homeostasis during aging. Levels of key ECM components are read out and communicated back into the cell to initiate a response that balances all the multifold components that make up the matrix. Interventions increasing key collagen levels are sufficient to drive enhanced remodeling during aging, whereas blocking this collagen enhancement leads to a collapse of this enhancement (Figure 3N). Furthermore, the enhancement of ECM is sufficient and required for cellular reprogramming upon longevity interventions.

### Screening identifies mechanotransduction genes as regulators for prolonged collagen expression

To investigate the above-described feedback loop and how longevity interventions prolong the maintenance of collagen dynamics, we designed a screen to assess ECM homeostasis during old age. Since the decline in the pattern I collagen transcription preceded its decline or remodeling out from the cuticle, we scored *col-144* promoter-driven GFP levels (P*col-144*::GFP) that progressively declined from day 1 to day 8 of adulthood (Figure 4A, Supplementary Figure 7A). At day 8 of adulthood, in a wild-type temperature-sensitive sterile background (*spe-9(hc88)*), most P*col-144*::GFP levels vanished, whereas long-lived mutants *glp-1(e2141)* or *daf-2(e1370)* still retained low levels of P*col-144*::GFP fluorescence, indicating prolonged collagen expression (Figure 4A). We aimed to identify two types of regulators by RNAi screening. The first, are negative regulators that would lead to higher P*col-144*::GFP levels at day 8 of adulthood upon knocking down by RNAi in a wild-type background, suggesting prolonged collagen maintenance as observed by longevity interventions. The second is for regulators required to prolong collagen expression in longevity-promoting mutant backgrounds, *i.e.,* RNAi knockdown would abolish higher P*col-144*::GFP levels at day-8 of adulthood in long-lived animals.

**Figure 4.**
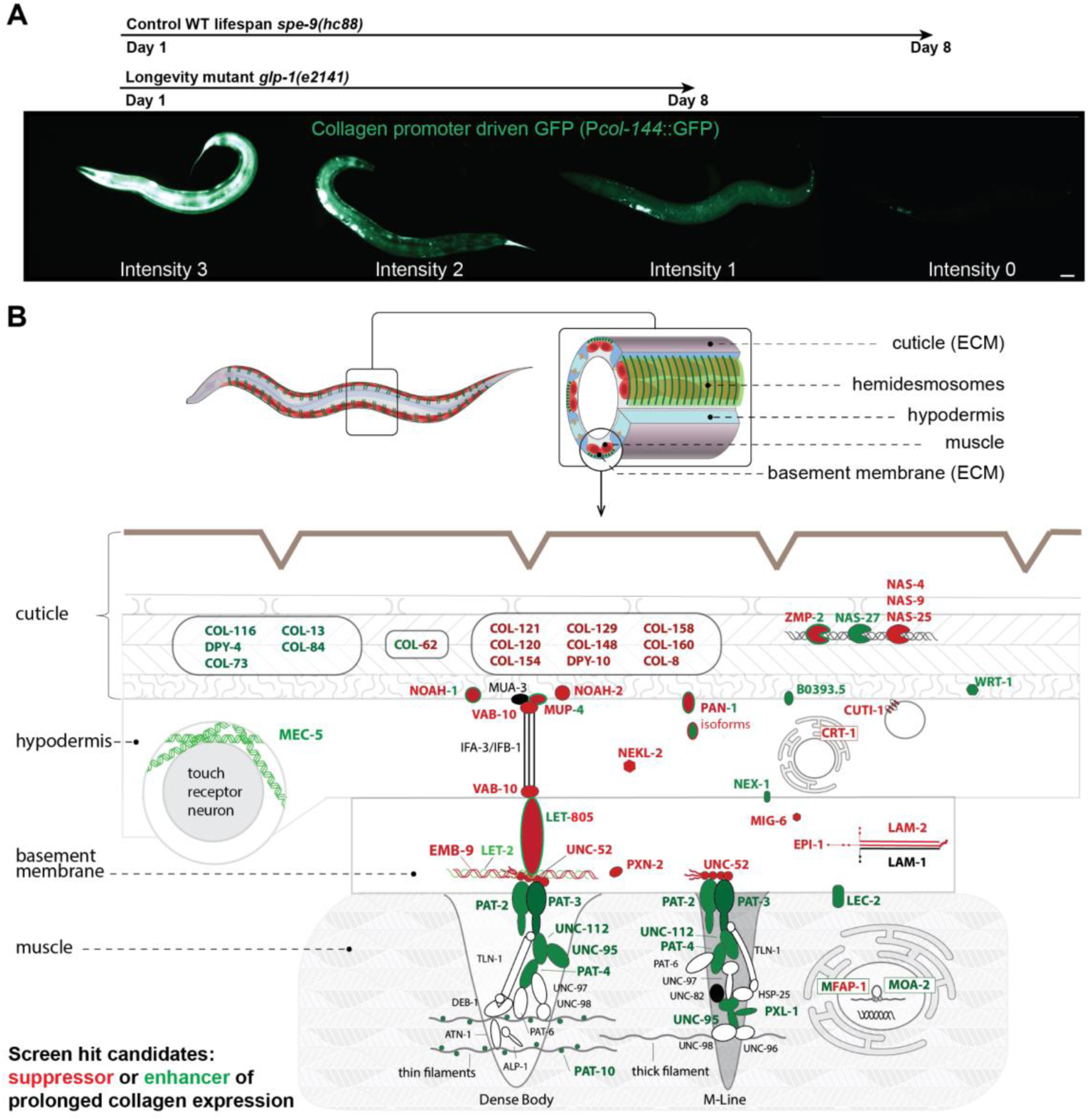
RNAi screen for transcriptional ECM regulators at old age. (A) Scoring scheme for the P*col-144*::GFP *C. elegans* RNAi screen. Above indicated with arrows are the usual decline of the fluorescence of the wild-type background *spe-9*(*hc88*) and long-lived mutant backgrounds *glp-1*(*e2141*). Scale bar = 50 µm. (B) The hits of the screen are shown in the predicted layers from *C. elegans* cuticle to body wall muscle (not drawn to scale). In green, RNAi hits led to higher P*col-144*::GFP at day 8 of adulthood. In red, RNAi hits that suppressed normally higher P*col-144*::GFP of long-lived mutants at day 8 of adulthood. The proteins in black were also tested but did not show a significant up- or down-regulation, while the proteins in white were not included in the screen. For details, see Supplementary Table 10.

We took a targeted approach to screen through most *C. elegans* kinases (382 out of 438 kinases), about one-third of all transcription factors (330 out of 934 genes), and 190 metabolism genes (Supplementary Figure 7B, Supplementary Table 10) (Venz et al., 2020). Since knockdown of certain collagens and ECM regulators abolishes prolonged collagen gene expression and longevity of normally long-lived *C. elegans* (Ewald et al., 2015), we generated a matrisome library containing 625 out of the 719 matrisome genes (Supplementary Figure 7B, Supplementary Table 10). Furthermore, a whole-genome RNAi screen has been previously performed for pathogen infections using a *col-12* promoter-driven reporter as a control (Zugasti et al., 2016). We re-analyzed their screening data for enhancers or suppressors of *col-12* expression during young adulthood and chose 133 genes (Supplementary Figure 7B, Supplementary Table 10). Lastly, we performed a literature search for ECM regulators across species and selected 98 genes. In total, we screened 1770 genes in more than three replicates and found 254 hits. Then in a second pass, we validated candidate genes in addition also in a *daf- 2(e1370)* longevity background and ended up with 107 confident hits (Supplementary Figure 7B, Supplementary Table 10).

We identified several required genes for longevity, such as *daf-16*/FOXO, *pqm-1,* and *xbp-1/*XBP1 (Supplementary Table 10) (Henis-Korenblit et al., 2010; Tepper et al., 2013). We also identified known longevity-promoting genes that, when knocked down. prolonged P*col-144*::GFP expression (*e.g., let-363/mTOR, sams-1/MAT1, pat-4/ILK1,* Supplementary Table 10) (Hansen et al., 2005; Kumsta et al., 2013; Vellai et al., 2003). Gene categories of candidate hits included autophagy, metabolism, molting, pathogen innate immune response, and signaling (Supplementary Figure 7C, Supplementary Table 10). To our surprise, most hits were in the matrisome and adhesome gene category (Supplementary Figure 7C, Supplementary Table 10). We mapped these candidates in an anatomical model displaying the four tissue layers: The cuticle (1) is attached to the hypodermis (2), which is attached to the basement membrane (3), which is attached to the body wall muscles (4; Figure 4B). We found that knocking down genes that form or remodel the cuticle either functions as an enhancer (green) or suppressor (red) of prolonged collagen expression (Figure 4B, Supplementary Table 10). RNAi of genes that anchor the hypodermis to the muscles (*mup-4/ matrilin, vab-10/ dystonin, let-805/ myotactin-fibronectin repeats*) via the basement membrane (*emb-9/collagen type IV, let- 2/ collagen type IV, unc-52/ perlecan, epi-1/ laminin alpha, lam-2/ laminin gamma*) were generally required for prolonged hypodermal collagen expression (Figure 4B, Supplementary Table 10). By contrast, knocking down genes that form the adhesome (*pat-2/ integrin alpha, pat-3/ integrin beta, unc-112/ pleckstrin, pat-4/ integrin-linked kinase, tln-1/ talin, pxl-1/ paxilin, deb-1/ vinculin, pat-10/* troponin C) upregulated and prolonged P*col-144*::GFP expression in the hypodermis (Figure 4B, Supplementary Table 10).

Most strikingly, all these genes are components that are either required for enabling or transmitting mechanical forces (mechanotransduction) during embryo elongation (Zhang et al., 2011). This suggests that the underlying mechanism of the feedback loop for prolonged collagen expression during aging might be mediated via mechanical force coupling across tissues.

### Progressive decline in colocalization of basement membrane components with adhesome during aging

For mechanotransduction to occur, both collagen type IV ([EMB-9]2 [LET-2]) and perlecan (UNC-52) bind to the integrin receptors composed of the heterodimers of integrin alpha INA-1 or PAT-2 with integrin beta PAT-3 (Figure 4B) (Gieseler, 2017; Teuscher et al., 2019a). To monitor the mechanical coupling of the basement membrane to integrin signaling during aging, we assessed the colocalization of collagen EMB-9 tagged with mCherry and integrin receptor beta PAT-3 tagged with GFP. Despite the age-dependent increase in EMB-9, UNC-52, and PAT-3 levels in muscular attachment structures, we found a progressive decline in colocalization of EMB-9 with PAT-3 during days 1 to 8 of adulthood, which was rescued by longevity intervention *daf-2(RNAi)* (Figure 5A-E, Supplementary Figure 8A-B, Supplementary Table 11).

**Figure 5.**
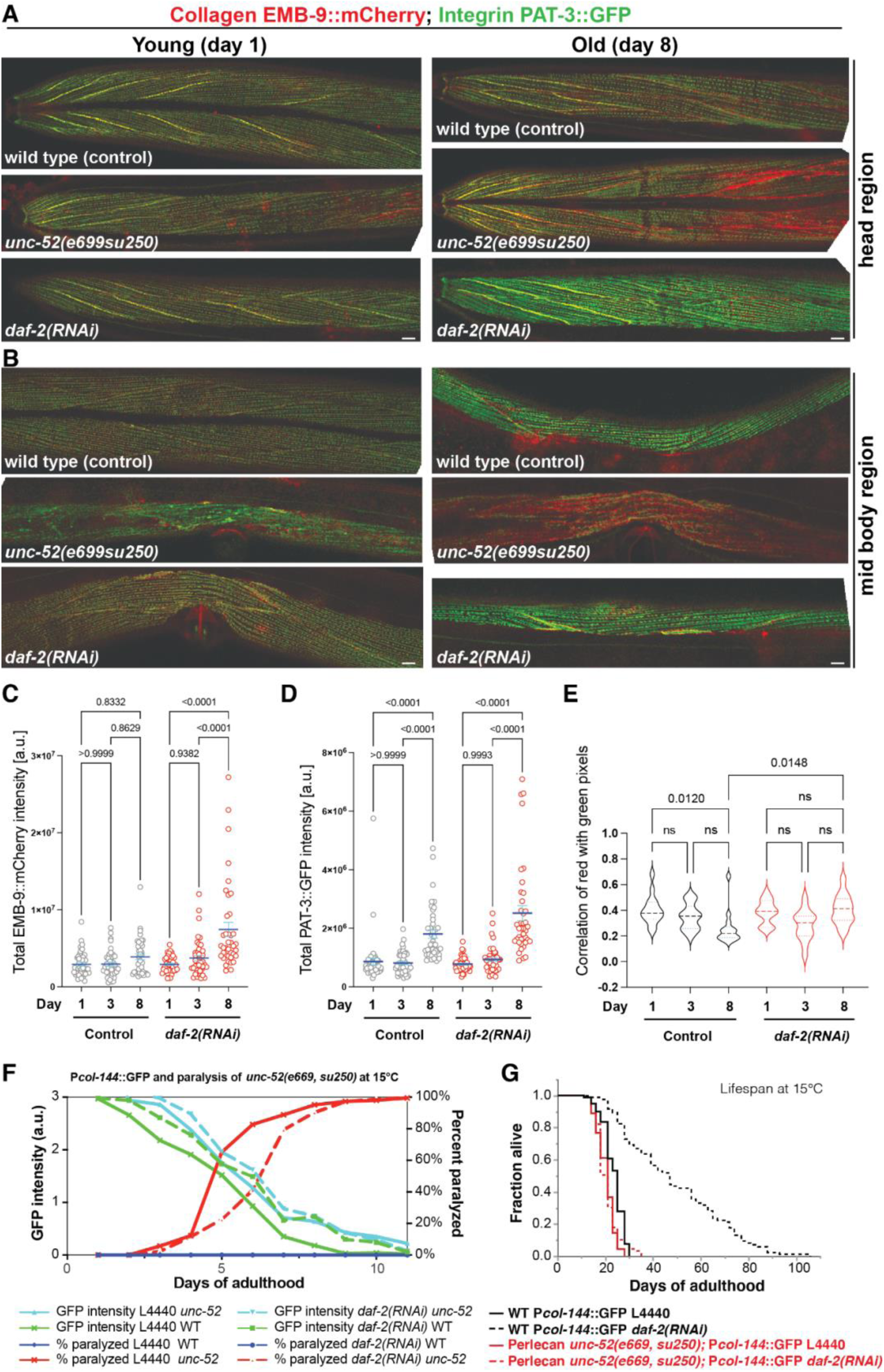
Disassociation of co-localization of collagen type IV from integrin β receptor. (A, B) Confocal image overlays of collagen type IV (EMB-9::mCherry) in red and integrin β receptor (PAT-3::GFP) in green of the head region (A) and midbody region (B) at day 1 and day 8 of adulthood. The yellow color indicates colocalization. Anterior to the left, ventral side down. Scale bar = 10 µm. (C, D) Quantification of total EMB-9::mCherry (C) and PAT-3::GFP (D) intensity levels. One-way ANOVA. (E) Quantification of colocalization by correlation of per-pixel red (EMB-9::mCherry) and green (PAT-3::GFP) intensities of the midbody region (B), which declined during aging in wild type but was maintained upon *daf-2(RNAi)* longevity intervention. One-way ANOVA. (A-E) For individual pictures, raw data, and statistical analysis, see Supplementary Table 11. (F) Longevity intervention *daf-2(RNAi)* prolonged collagen expression (P*col-144*::GFP) of wild type (green line) during aging but not in a perlecan *unc-52(e669, su250)* mutant background (aquamarine line) at permissive temperature 15°C. The age-dependent *unc- 52(e669, su250)* mutant paralysis phenotype (red line) was delayed by *daf-2(RNAi)* (dashed red line) at 15°C. For details, see Supplementary Table 12. (G) Continuing with these same animals, *unc-52(e669, su250)* mutants completely suppressed *daf-2(RNAi)* longevity at 15°C. For details, see Supplementary Table 7.

Because perlecan UNC-52 is at the interface between collagen EMB-9 and integrin receptor PAT-3, we used a temperature-sensitive perlecan *unc-52(e699, su250)* at the semi-permissive temperature of 20°C and found that loss of *unc-52* function accelerated the loss of colocalization of EMB-9 form PAT-3 during day 1 to day 8 of adulthood (Figure 5A, 5B), presumably leading to detachment of muscle from the basement membrane. This is consistent with the observation that these *unc-52(e699, su250)* animals become progressively paralyzed in the midbody region at permissive temperature (15°C) during aging (Ben-Zvi et al., 2009) but not in the head region. coinciding with the increased colocalization observed in the posterior head region (Figure 5A).

### Mechanical coupling is required for longevity

To understand the relationship between loss of mechanical tension across tissues and longevity, we used the perlecan *unc-52(e699, su250)* paralysis phenotype as a functional read-out and hypodermal collagen reporter (P*col-144*::GFP) as a read-out for the loss of mechanical coupling to gene expression across tissues. First, we noticed that the perlecan *unc-52(e699, su250)* had higher base-line P*col-144*::GFP expression during development and throughout adulthood (Supplement Figure 8C-D, Supplementary Table 12), suggesting that the strength of mechanical coupling determines gene expression levels across tissues, which is consistent with a feedback loop to adapt exoskeleton cuticle strength with muscle strength.

We found that *daf-2(RNAi)* postponed the perlecan *unc-52(e699, su250)* paralysis phenotype during adulthood by two days at a permissive temperature of 15°C and a semi-permissive temperature of 20°C (Figure 5F, Supplementary Figure 8E). This is consistent with a previous study showing a delay of *unc-52*-paralysis in long-lived *glp-1* mutants (Cohen-Berkman et al., 2020), suggesting that this delay in paralysis is due to the general improvement of protein homeostasis promoted by longevity interventions. By contrast, the prolonged collagen gene expression and extreme longevity upon *daf-2(RNAi)* were blunted and completely abolished by perlecan *unc-52(e699, su250)* mutations at 15°C or 20°C, respectively (Figure 5F, 5G, Supplementary Figure 8E, Supplementary Table 7, 12). This demonstrates that proper tissue coupling is required to promote hypodermal collagen expression for the systemic longevity effects.

### Relevance of the matrisome and adhesome for longevity

To identify which of these components are functionally important for longevity, we measured the lifespan of 35’795 individuals, including 39 matrisome and adhesome mutants treated either with control or *daf-2(RNAi)* (Figure 6A, Supplementary Table 7) using the lifespan machine (Stroustrup et al., 2013). We found requirements for *daf-2*- longevity across the different matrisome categories (Supplementary Figure 9), demonstrating the essential interplay of these molecular components to form a proper functional network. In line with our screening data, components of both ECMs, basement membrane and cuticle, were required (Figure 6A). Consistent with our proteomics data, whereby the age-dependent decline of perlecan/UNC-52 protein levels was counteracted by *daf-2(RNAi)* (Figures 1 and 3), these *unc-52* mutants showed the strongest epigenetic requirements for *daf-2*-longevity (Figure 6A).

**Figure 6.**
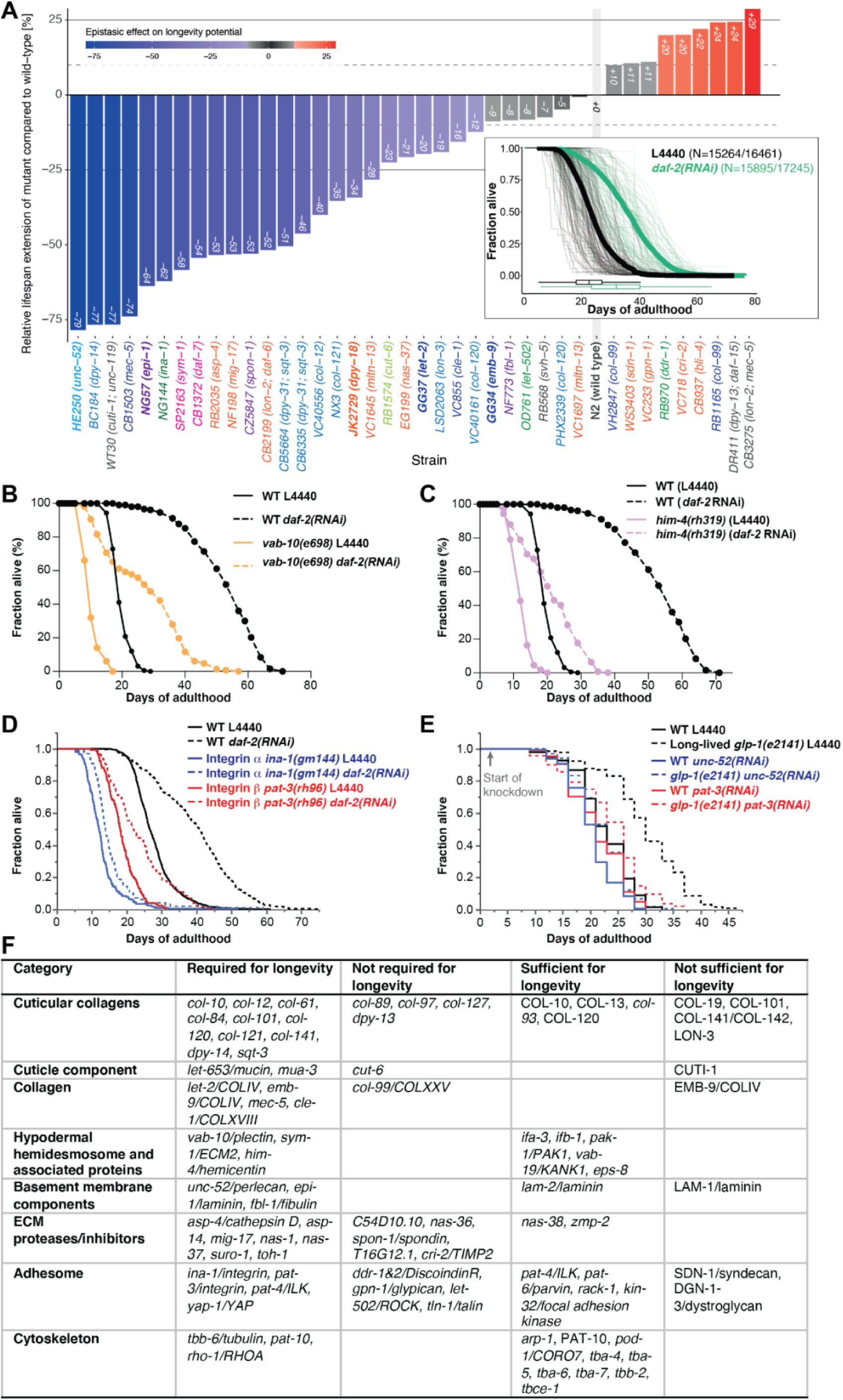
Functional matrisome is of key importance for *C. elegans* longevity. (A) Epistatic effect of matrisome mutations on rIIS-mediated longevity potential. Bars represent the relative differences in mean lifespan extension measured in matrisome mutants compared to wild-type animals within the same batch. (B) Mutation in *vab-10*/plectin/dystonin shortened wild type (WT) lifespan and blocked longevity upon reduced insulin/IGF-1 receptor signaling at 20°C. (C) Mutation in *him-4*/hemicentin shortened wild type (WT) lifespan and blocked longevity upon reduced insulin/IGF-1 receptor signaling at 20°C. (D) Mutations in integrin **α** and β suppressed *daf-2(RNAi)* longevity at 20°C. (E) Knocking down perlecan/*unc-52* or integrin β/*pat-3* starting at day 2 of adulthood suppressed germ cell-less (*glp-1(e2141)*) mediated longevity. (F) Summary of all matrisome and adhesome genes implicated in longevity. RNAi or genetic mutants are in italics, overexpression is in capital letters. (A-F) For details, statistics, and additional trials, see Supplementary Table 7.

Next, we wondered whether the hemidesmosome components discovered in our screen would affect lifespan. We found that *vab-10/*plectin loss-of-function mutants were shorter-lived and blocked *daf-2*-longevity (Figure 6B, Supplementary Table 7). Hemicentin *him-4* is essential for hemidesmosome anchoring and mechanotransduction (Vogel and Hedgecock, 2001), and *him-4* was required for longevity (Figure 6C, Supplementary Table 7).

Since hemidesmosomes interact with integrin receptors (Gieseler, 2017), we next investigated the integrin receptors’ requirements for longevity. Both. integrin alpha (*ina- 1*) and the sole integrin beta (*pat-3*) were required for *daf-2-*longevity (Figure 6D, Supplementary Table 7). To exclude any developmental effects and to test another longevity pathway, we used long-lived *glp-1(e2141)* animals and knocked down *unc-52* or *pat-3* by RNAi starting at day 2 of adulthood. Adulthood-specific knockdown of *unc-52* or *pat-3* had no lifespan effects on normal-lived control animals but abolished the longevity of *glp-1(e2141)* mutants (Figure 6E, Supplementary Table 7). Thus, integrin receptors are required for longevity but not for wild-type aging, consistent with the observation that only longevity interventions enhance pattern I collagen expression. This implicates a functional role of mechanotransductive integrin signaling in this feedback loop to enhance extracellular matrices.

Integrins transmit mechanical forces from the ECM to the cytoskeleton, and vice versa, whereby they get physically connected to the cytoskeleton by linker proteins, including talin/TLN-1 and integrin-linked kinase (ILK/PAT-4), and RHO-associated kinase (ROCK/LET-502) regulate the dynamic reorganization of cytoskeletal proteins by regulating the phosphorylation of myosin motors (Humphrey et al., 2014). Adult-specific upshift to 20°C of temperature-sensitive loss-of-function ROCK/*let-502* mutants did not suppress *daf-2(RNAi)* longevity (Supplementary Figure 10A, Supplementary Table 7). While the dephosphorylation of the integrin receptors at tyrosine 792 in the membrane-proximal NPXY motif promotes integrin activations via talin recruitment (Walser et al., 2017), phospho-deficient integrin PAT-3(Y792F) mutants, as well as talin/*tln-1* mutants treated with *daf-2(RNAi)*, were still long-lived (Supplementary Figure 10B, Supplementary Table 7). Loss of focal adhesion kinase *kin-32* increased lifespan (Supplementary Figure 10C, Supplementary Table 7). Heterozygous ILK/*pat-4* mutants were shorter-lived and blocked *daf-2*-longevity (Supplementary Figure 10D, Supplementary Table 7), consistent with severe *pat-4* knockdown leads to detachment of the cytoskeleton and shortening of lifespan, whereas mild *pat-4* knockdown has a mild effect on cytoskeleton detachment and increases lifespan (Nishimura et al., 2014). Consistent with our screening hits and proteomics data are cytoskeleton remodelers implicated in longevity by our and other groups (Figure 5F) (Baird et al., 2014; Ewald et al., 2017; Mansfeld et al., 2015; Vitiello et al., 2021). This points towards a hemidesmosome-to-integrin-to-cytoskeleton remodeling axis to mediate downstream mechanotransduction and organismal longevity.

### Yap-1 is required for longevity and collagen homeostasis

To determine the downstream effectors of this hemidesmosome-to-integrin-to-cytoskeleton remodeling axis, we explored whether YAP is important for longevity, as the conserved Yes-associated protein transcriptional co-activator YAP is implicated in the transcriptional response to ECM stiffness and cytoskeletal organization (Dupont et al., 2011; Panciera et al., 2017). Treatment of *yap-1(tm1416)* putative null mutants with *daf- 2(RNAi)* revealed that the loss of *yap-1* function abolished longevity upon reduced Insulin/IGF-1 signaling (Figure 7A, Supplementary Table 7). We did not observe any enhanced YAP-1 nuclear localization upon *daf-2(RNAi)* but observed higher levels of YAP-1, which was potentiated at 25°C (Figure 7B, 7C, Supplementary Table 13). Interestingly, under normal conditions, very faint YAP-1::GFP signal was observed at hemidesmosome-containing structures (Figure 7D-H, Supplementary Table 13). Upon *daf-2(RNAi)*, YAP-1 translocated from the cytoplasm to localize at the hemidesmosome- containing structures between the apical and basal VAB-10/plectin/dystonin (Figure 7D- H, Supplementary Table 13). This might suggest that YAP-1 could help read out mechanical changes occurring at these hemidesmosome-containing structures. In mammals, YAP responds to a broad range of mechanical cues, from shear stress to cell shape, and extracellular matrix rigidity, and is considered a mechano-sensitive transcriptional regulator (Elosegui-Artola et al., 2017; Moya and Halder, 2019; Panciera et al., 2017). To test YAP-1’s mechano-sensitive role in *C. elegans*, we placed L4 animals for 3 days under ca 12 Pa pressure (Supplementary Table 13). We found that YAP-1 levels increased under these mildly higher pressure conditions (Figure 7I, Supplementary Table 13), suggesting its expression responded to mechanical compression. If our model of hemidesmosomes regulating collagen expression via mechanical tensions is correct, then placing *C. elegans* under these mild pressure conditions should prolong collagen expression during aging. To control for general gene expression changes under pressure, we generated *C. elegans* expressing mCherry in body wall muscles and neonGreen in the hypodermis under the *col-120* promoter, whose expression rapidly declined in early adulthood (Figure 7J). Normalized to control muscular mCherry expression, which did not change under pressure, we found that *col-120* expression was enhanced by these 3 days of mild pressure (Figure 7K, Supplementary Table 14). We confirmed this enhanced collagen expression under pressure by *col-144* promoter-driven GFP in a sterile background avoiding FUdR (Figure 7L, Supplementary Table 14). Weakening the hemidesmosome force transduction by *unc-52* temperature-sensitive mutations, as before, showed already higher *col-144* expression under normal conditions but blunted the enhancement of collagen expression upon pressure (Figure 7M, Supplementary Table 14). Furthermore, knockdown of *yap-1* abolished this pressure-mediated collagen expression (Figure 7O, Supplementary Table 14), suggesting that YAP-1 requirements for longevity are due to responding to mechanical tension changes from hemidesmosomes to coordinate collagen expression (Figure 7P).

**Figure 7.**
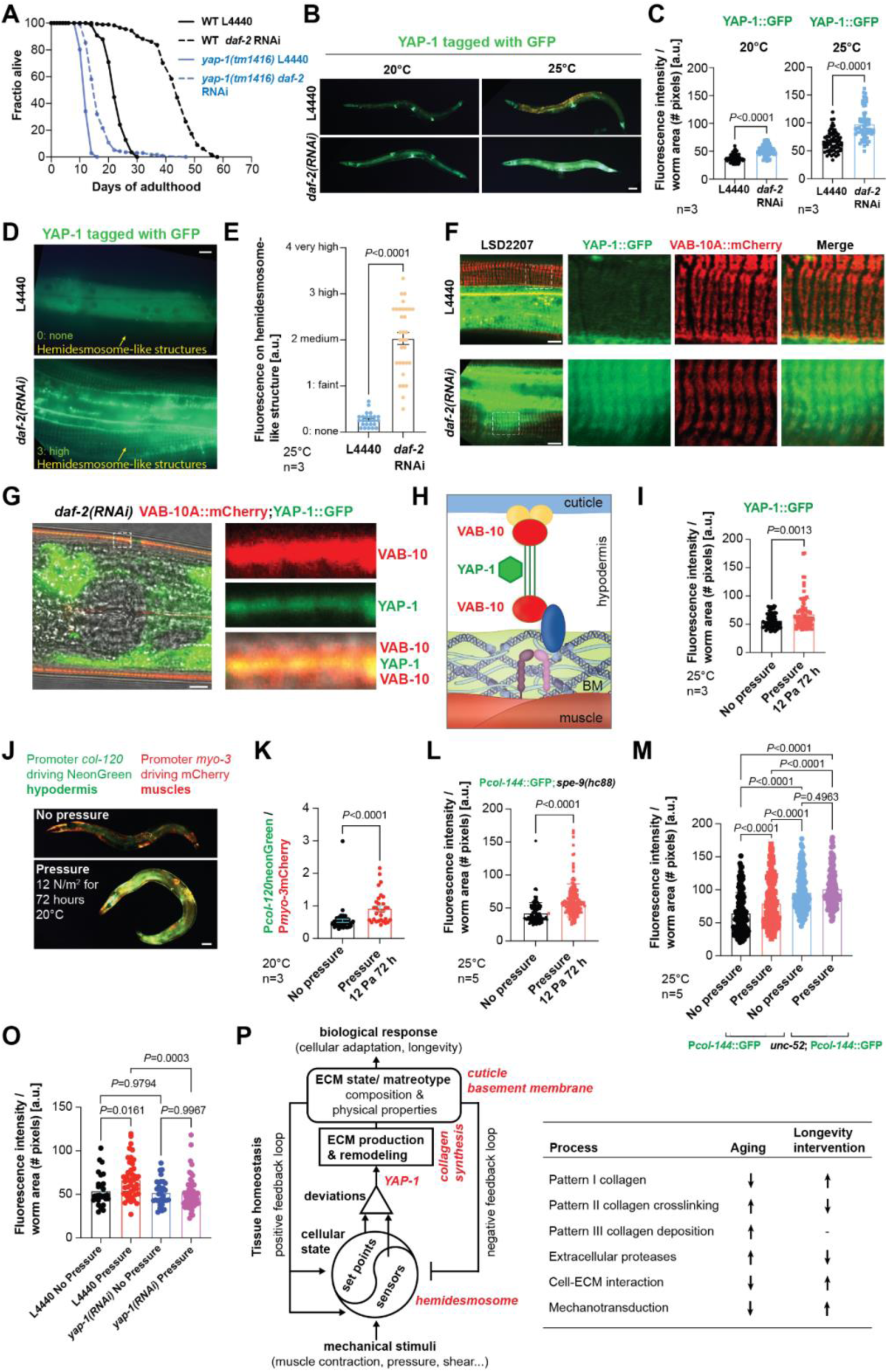
YAP-1 is required for longevity and pressure-induced collagen expression during aging. (A) Loss-of-function mutants of *yap-1(tm1416)* abolish longevity upon reduced Insulin/IGF-1 signaling. For raw data, additional trials, and statistics, see Supplementary Table 7. (B-C) YAP-1::GFP expression was increased upon reducing insulin signaling. (B) Representative images show increased expression of *ihIs35* YAP-1::GFP in animals fed with *daf-2* RNAi as compared to L4440 control at 20°C and 25°C. Scale bar = 50 µm. The graph shows the quantified data for the YAP-1::GFP expression. 3 independent biological trials. Welch’s *t*-test was used for significance analysis. For raw data and statistical details, see Supplementary Table 13. (D-E) YAP-1::GFP expression localizes at hemidesmosome-containing structures was increased upon reducing insulin signaling at 25°C. (D) Representative images show increased expression of *ihIs35* YAP-1::GFP in animals fed with *daf-2* RNAi as compared to L4440 control at 25°C. Scale bar = 10 µm. (E) The graph shows the distribution of the categories of hemidesmosome-containing structures. 3 independent biological trials. Welch’s *t*-test was used for significance analysis. For raw data and statistical details, see Supplementary Table 13. (F-G) Upon *daf-2(RNAi)* at 25°C, more YAP-1 was colocalized with VAB-10. The boxed areas are enlarged on the right showing the *ihIs35* YAP-1::GFP (green channel), *mc100* VAB-10A::mCherry (red channel), and merge of the confocal images using the transgenic strain LSD2207. Scale bar = 10 µm. For additional images and individuals, see Supplementary Table 13. (H) Schematic of YAP-1 localizing at the hemidesmosome-containing structure between the apical and basal VAB-10 based on confocal images shown in (G). Not drawn to scale. (I) YAP-1::GFP expression was increased upon constant pressure for three days at 25°C. 3 independent biological trials. Welch’s *t*-test was used for significance analysis. For raw data and statistical details, see Supplementary Table 13. (J-K) Pressure induces collagen expression in the hypodermis. (J) Representative images of LSD1126 P*col-120*::mNeonGreen; P*myo-3*::mCherry transgenic *C. elegans* either kept under 12 Pa pressure for 72 h starting at L4 at 20°C or not. Scale bar = 50 µm (K) Quantification of 3 independent biological trials. A *t*-test was used for significance analysis. For raw data and statistical details, see Supplementary Table 14. (L) Promoter-driven *col-144*::GFP expression in a temperature-sterile background (LSD2002) was increased upon constant 12 Pa pressure for three days at 25°C. 5 independent biological trials. Welch’s t-test was used for significance analysis. For raw data, additional trials also at 20°C, and statistical details, see Supplementary Table 14. (M) Mutations in *unc-52(e669,su250)* blunted promoter-driven *col-144*::GFP expression upon constant 12 Pa pressure for three days at 25°C. 5 independent biological trials. One- way ANOVA was used for significance analysis. For data and statistical details, see Supplementary Table 14. (O) Knockdown of *yap-1* abolished the promoter-driven *col-144*::GFP expression upon constant 12 Pa pressure for three days at 25°C. 1 independent biological trial is shown. One-way ANOVA was used for significance analysis. For data, additional trials, and statistical details, see Supplementary Table 14. (P) Proposed biomechanical model of ECM homeostasis and longevity. The right model is adapted for our *C. elegans* finding from mechanobiological regulation of arterial walls, which include smooth muscles, endothelial cells, fibroblasts, and ECM (by Humphrey and Schwartz 2021). The left model depicts the ECM homeostasis-related process that declines during aging but is counteracted by longevity interventions.

## Discussion

There is a general appreciation of the importance of the progressive decline of ECM integrity during aging (Ewald, 2020). However, much of the past analysis focused on physical and structural disorganization, which is improved by longevity interventions (Essmann et al., 2020; Rahimi et al., 2022). Our previous genetic study pointed toward the importance of ECM components at the molecular level for promoting longevity (Ewald et al., 2015). Through the establishment of matreotypes using proteomics and *in-vivo* reporter systems, we have laid here the groundwork for understanding a number of distinct features of the dynamic ECM composition changes during aging and longevity.

Because of the long a half-life time of 114 years of collagens in humans (Ewald, 2020) it is generally thought that when ECMs are formed in development, collagens would simply serve as an inert scaffold and accumulate damage during aging, such as collagen fragmentation through dysregulated ECM proteases and accumulation of AGEs leading to crosslinking and stiffening of ECMs (Ewald, 2020). While we observed an age- dependent increase in proteases which was reverted by longevity interventions, we additionally characterized three distinct patterns of collagen dynamics resembling changes occurring during mammalian aging and diseases (loss of collagen mass, collagen crosslinking, and localized collagen deposition). Consistent with the hypergrowth or “faucet left on” theory (Blagosklonny, 2012), we found that *C. elegans* pattern III collagens are continuously synthesized and integrated into the basement membranes analogous to the increase of human basement membrane thickness during aging (Halfter et al., 2015). By contrast, *C. elegans* pattern II collagens are synthesized once and integrated into the ECM, and they thus suffer age-related consequences (*i.e.*, crosslinking and stiffening of ECM, analogous to changes of mammalian or human tendons (Haefke and Ewald, 2020; Hamlin and Kohn, 1972). Interestingly, *C. elegans* pattern I collagens are the most dynamic collagens, which become excised out of the ECM during aging but are replenished for longer periods under longevity-promoting conditions. This is analogous to the human skin during aging, whereby collagen mass progressively declines (1% per year), and loss of mechanical tension-coupling results in declined collagen synthesis (Shuster et al., 1975; Varani et al., 2006). Another parallel is the observation that certain collagens are overexpressed in centenarians based on proteomic signatures (Sebastiani et al., 2021). Most importantly, enhancing *C. elegans* pattern I collagen dynamics is a prerequisite for longevity. We identified a regulatory feedback loop reading out the key pattern I collagens to signal into the cells to not only adjust ECM components to form a functional matrix but also to reprogram metabolism, stress defense, and cellular homeostasis upon longevity interventions (Figure 7P). We undertook three different approaches (proteomics, lifespan assays, genetic screening) to identify the underlying molecular mechanism(s) of this essential-for-longevity feedback loop. From all the imaginable possibilities, surprisingly, all three approaches pointed to one molecular hemidesmosome-containing structure spanning from the cuticle through the hypodermis, through the basement membrane to the muscles.

Although not formally implicated, there are several genetic observations during development that point to a reciprocal role of hemidesmosome structures in mechanosensation and ECM remodeling. The hemidesmosome-containing structures are required for contractile force transmission for normal locomotion of *C. elegans (Hresko et al., 1999).* Muscle contractions are essential for embryonic elongation by mechanical coupling and proper assembly of these filamentous hemidesmosome- containing structures from the muscle basement membrane through the hypodermis to the outer cuticle that acts as a soft exoskeleton (Figure 4B) (Zhang et al., 2011). Muscular contraction induces mechanical force transmission promoting reorganization of actin bundles in the hypodermis (Lardennois et al., 2019), indicating a mechanotransductive remodeling adapting inner and outer tensions and forces to these scaffolding structures.

Besides force transmission and cytoskeleton adaptation, genes that influence the physical properties of ECM were also identified in our screen during aging. For instance, *noah-1* and *noah-2* are important for maintaining mechanoreceptor potentials and cuticular ECM remodeling (Frand et al., 2005; Vuong-Brender et al., 2017). Intriguingly, loss of *unc-95*, *unc-52,* or *pat-3* results in ECM-associated molting defects (Frand et al., 2005; Zaidel-Bar et al., 2010). By contrast, mutations in muscle myosin *unc-54* important for muscle contraction or mutations in *unc-13* important for neurotransmitter release did not induce cuticular collagen expression (Broday et al., 2007). Moreover, scanning electron microscopy revealed abnormal, branched, or flat cuticular annuli of *unc- 52*/perlcan mutants, but not for *unc-13*/neurotransmitter-release mutants (Broday et al., 2007), suggesting that impaired hemidesmosome function impairs proper cuticle ECM morphology. In our screen, we also identified *pxn-2*/peroxidasin that promotes sulfilimine crosslinks of basement membrane type IV collagen, thereby regulating basement membrane mechanical properties. Defects in *pxn-2* can be bypassed by mutations in hemidesmosome components, such as *let-805, vab-10,* and *unc-52* (Gotenstein et al., 2018), suggesting changes in either the cuticular or basement membrane ECM are mediated by hemidesmosomes.

Mechanical compression of hemidesmosomes by placing *C. elegans* into hypergravity impairs the migration of motor neurons over the muscle, and mutations in *vab-10*, *unc-52,* and other hemidesmosome components rescue neuronal migration (Kalichamy et al., 2020). Vice versa, stretching hemidesmosomes unmasks the SH3 domain of VAB-10/Plectin enabling mechanosensitive signaling essential for embryonic elongation (Suman et al., 2019). Interestingly, hydrostatic pressure increases *col-107* mRNA and increases lifespan (Watanabe et al., 2020). Similarly, a three-month space flight hypogravity induces collagen turnover in mice’s skin (Neutelings et al., 2015).

Together with our results that exogenously applied pressure promotes collagen expression and longevity, we propose that the physical properties of either the cuticle ECM or basement membrane ECM are sensed and adapted by hemidesmosomes to orchestrate force coupling across tissues (Figure 7P). Our proposed feedback loop of ECM homeostasis via hemidesmosome tension coupling reconciles and explains these previous observations during development (Figure 7P). Moreover, we adapted a biomechanical model recently established by Humphrey and Schwartz describing arterial wall integrity, where mechanical forces regulate gene expression and signaling across several cell types, smooth muscle cells, endothelial cells, fibroblasts, and surrounding ECM to maintain optimal functions (Humphrey and Schwartz, 2021), which captures well our findings. Thus, *C. elegans* recalibrates tissue forces applied to ECM through remodeling ECM composition (Figure 7P).

The key question is how do changes in hemidesmosomes affect longevity? During aging, we observed uncoupling of basement-membrane collagen type IV from its integrin receptor, suggesting an age-dependent loss of hemidesmosome integrity. In early adulthood of *C. elegans,* the protein homeostasis network collapses, which is slowed by longevity interventions (Ben-Zvi et al., 2009; Labbadia and Morimoto, 2014). Similarly, we found that longevity interventions were able to slow this age-dependent uncoupling of ECM from its receptor (Figure 5). We favor the idea that loss of proteostasis drives the loss of hemidesmosome tension-coupling, since a less stable perlecan mutant, serves as a readout of ECM proteostasis (Ben-Zvi et al., 2009), accelerated this disassociation (Figure 5). Mutated perlecan disorganizes and forms aggregates, which are removed by enhanced chaperones, autophagy, and proteasomal functions (*i.e.,* better protein homeostasis) of longevity mutants or drugs (Alavez et al., 2011). Furthermore, in our screen, we identified calreticulin *crt-1*/CALR, which regulates UNC-52/perlecan folding and protein levels (Zahreddine et al., 2010). Thus, longevity interventions might improve the integrity of hemidesmosomes to ensure their dynamic adaptation and signaling.

Beyond the known fact that ECM-cell detachment leads to apoptosis or loss of cellular identity (He et al., 2015), several recent findings point toward the idea that hemidesmosomes might directly regulate cellular homeostasis and tissue adaptation. For instance, chaperone HSP-43 is constitutively expressed and stored at hemidesmosomes (Fu et al., 2020). Upon heat, HSP-43 is released from the hemidesmosomes as a fast response, analogous to small heat shock chaperones associated with desmosomes and focal adhesions to ensure resistance against heat-induced damage in mammalian cells (Fu et al., 2020). Disturbances or disintegration of hemidesmosomes induces V-ATPase and activates lysosomes to facilitate ECM turnover during development (Miao et al., 2020). Hemidesmosomal integrity is also linked to mitochondrial ATP production and muscle protein synthesis (Etheridge et al., 2015). Moreover, mitochondrial homeostasis and stress responses (mitohormesis) are interlinked with ECM-integrin cytoskeleton remodeling in *C. elegans* and human stem cells (Ding et al., 2008; Munkácsy et al., 2016; Schinzel et al., 2019; Tharp et al., 2021). Lastly, disruption of the cuticle or the upper part of the hemidesmosomes (*mup-4*) induces a pathogen response (Zhang et al., 2015). Similarly, in primary human epidermal keratinocytes, disruption of hemidesmosomes induces an antimicrobial peptide pathogen response (Zhang et al., 2015). Intriguingly, our data shows that the hemidesmosome feedback loop that enhances collagen expression leads to cellular adaptation of cytoskeleton, metabolic, oxidative stress response, and pathogen response proteins (Figure 7P, Supplementary Table 9), all processes important for longevity interventions to maintain cellular homeostasis.

Mechanotransduction refers to the conversion of biophysical forces into transcriptional output or adaptation. Our lifespan screening identified mechano- responsive transcriptional co-activator YAP-1, which is required for longevity. Consistent with our proteomics on downstream processes upon enhanced collagen overexpression, YAP-1 is required for cytoskeleton, stress, and pathogen responses (Iwasa et al., 2013; Lee et al., 2019; Ma et al., 2020). Our data showed that YAP-1 levels are increased at higher temperatures and YAP-1 is found at the hemidesmosome-containing structures under reduced insulin/IGF-1 signaling. Furthermore, we demonstrated that YAP-1 responds to externally applied pressure and promotes the enhancement of collagen expression requiring intact hemidesmosomes (Figure 7). In humans, YAP1 senses the stiffness of the ECM (Elosegui-Artola et al., 2017), which increases with age leading to dysregulation of YAP1 impairing mechanotransduction and altering stem cell differentiation *in vivo* (Pelissier et al., 2014) and inducing cellular senescence *in vitro* (He et al., 2019; Xie et al., 2013). A feedback loop of YAP regulating collagen deposition and ECM remodeling has been observed for cancer-associated fibroblasts (Calvo et al., 2013). Although activation of YAP1 promotes tissue regeneration, hyperactivated YAP1 has been observed in human cancers, suggesting a tight regulation of YAP1 activity for regenerative medicine (Moya and Halder, 2019). Interestingly, several phenotypes we observed here are conserved in mammals, for instance, YAP1 is involved in heat stress response (Luo et al., 2020), cytoskeletal dynamics (Morikawa et al., 2015), pathogen response (Wang et al., 2017), and is implicated in hemidesmosome-structure related human epidermolysis bullosa disease (Rosa et al., 2019). However, to the best of our knowledge, our results are the first direct non-invasive *in-vivo* data showing the mechanotransduction of YAP-1 on collagen expression and linking its action to hemidesmosomes as a feedback homeostatic regulator.

The hemidesmosome-containing structure identified here that is important for longevity might be an ancestral structure combining two conserved mechano-regulators, whereby the upper part resembles hemidesmosomes found in mechanical exposed epithelial tissue (*e.g.,* skin, esophagus, intestine) and the lower part, from the basement membrane to the muscle, resembles focal adhesions (Figure 4B). In mammals, there is dynamic crosstalk between hemidesmosomes and focal adhesions. For instance, during cell migration and wound healing, hemidesmosomes cluster as ordered arrays interspersed by actin-associated focal adhesions (Hatzfeld and Magin, 2019; Molder et al., 2021). Furthermore, cellular tension induces the dynamic interplay between hemidesmosomes and focal adhesions leading to the activation of YAP *in vitro* (Wang et al., 2020), suggesting a functionally conserved mechanism.

Given this conserved interplay, impairment of these structures has several consequences for mammalian aging reminiscent of our findings. Facial skin biopsies from older humans (>60 years) show disassociation of ꞵ4-integrin from hemidesmosomes and decreased collagen type IV levels compared to younger individuals (Varlet et al., 1998).

Furthermore, in human skin, an age-dependent decline of collagen COL17A1 which is part of hemidesmosomes is observed, which drives the loss of stem cell maintenance and skin aging in mice (Liu et al., 2019). A recent clinical trial (NCT03536143) has shown that it is feasible and safe to administer hemidesmosome components by topical viral application in recessive dystrophic epidermolysis bullosa (Gurevich et al., 2022), suggesting that targeting the age-dependent decline of hemidesmosomes might be possible as a therapeutic intervention. The second line of convergent consequences of losing the lower part of the hemidesmosomes is the commonly observed detachment of muscle from the basement membrane via loss of focal adhesions and other receptors that might drive muscular dystrophy (Yurchenco et al., 2004). Muscular dystrophy patients die in their early twenties (Broomfield et al., 2021) and also show muscular fibrosis (Zhou and Lu, 2010), *i.e.*, excessive collagen production that can be *ex vivo* and *in vitro* reproduced by either a stiffer environment or more contractile cytoskeleton (Engler et al., 2004; Griffin et al., 2005), hinting at a derailed outside-in or inside-out feedback loop. Interestingly, a previous *in-silico* study found focal adhesions as a central network associated with human longevity (Wolfson et al., 2009). Thus, this suggests a conserved role of mechanical forces on focal adhesion and hemidesmosomes in aging and longevity.

In summary, although mechanotransduction has been observed to derail with aging, we provide here the first *in-vivo* evidence that a hemidesmosome feedback loop is essential to promote organismal lifespan extension. We found coordination between two distinct ECM adjusting ECM remodeling across tissues. We demonstrated that during aging, this ECM remodeling declines, and longevity interventions maintain this ECM remodeling, which promotes tissue integrity and cellular adaptations. Identifying mechanotransduction and hemidesmosomes as novel targets promoting longevity might open new therapeutic avenues.

## Limitations of the study

While our methodology and results postulate a new model and are a significant step forward towards elucidating the protein homeostasis and remodeling of ECMs elicited by longevity interventions *in-vivo* in an organism, the study also has a number of limitations. First, although we identified many ECM proteins reproducibly, due to inherent technical challenges of solubilizing the inherently poorly soluble ECM fraction for proteomics, this data is likely incomplete for at least some ECM proteins at some time points during the longitudinal time course. Second, we used multicopy translational reporters of cuticular collagens, since the collagens of interest, when endogenously tagged with CRISPR, showed too weak of a fluorescent signal in the ECM, which was masked by the age-related autofluorescence. Third, collagens are extracellularly cleaved and crosslinked in the ECM and several attempts of GFP tagging collagens, including between triple helices, disrupted matrix morphology. This warrants for future development of smaller tags and specialized microscopy to overcome these challenges. Forth, while our study covered a representative collection of core-matrisome proteins (*e.g.,* 3 out of 35 ECM glycoproteins, 1 out of 3 proteoglycans, 4 out of 7 conserved collagens, 11 out of 174 cuticular collagens, 3 out of 3 integrins, and 2 out of 2 hemidesmosome-like receptors), limited by the above-mentioned technical challenges, a more complete assessment of ECM components could further strengthen our conclusions. Fifth, we have shown an age-dependent decline in colocalization of basement-membrane collagen with integrin receptors. Although, we and others (Alavez et al., 2011; Ben-Zvi et al., 2009; Cohen-Berkman et al., 2020) demonstrated that genetically disrupting this interaction leads to muscle detachment from the basement membrane and a paralysis phenotype, proving physical disassociation turned out to be challenging, either due to difficulties in the preparation of the insoluble collagens or due to the fact that integrins bind collagens via a catch-bond mechanism (Kong et al., 2009), and warrants further investigation. Lastly, while previous isolated studies have shown that several of our key findings are also relevant in higher organisms, an experimental investigation and validation of the discovered mechanisms in mammals will be key for future translation of our findings for therapeutic purposes.

## Author contributions

All authors participated in analyzing and interpreting the data. CYE, ACT, and CS designed the experiments. CS, AG, AMH, SAD, and CYE performed lifespan assays. AMH and CS performed the comparative automated lifespan analysis. CS and AF performed proteomics. SAD performed FRET-sensor experiments. IS programmed FRET image analysis. CS performed the bioinformatic analysis. MH quantified cuticle thickness. OG programmed and performed the image analysis. AG and CYE performed the pressure experiments. ACT performed all other experiments. CYE wrote the manuscript in consultation with the other authors.

## Author Information

The authors have no competing interests to declare. Correspondence should be addressed to C. Y. E.

## Acknowledgment

We thank Mariam Baghdady, Kieran Toms, Vira Chea, and Katharina Tarnutzer for help with the screen, Elisabeth Jongsma for her help scoring GFP, Özlem Altintas for help with lifespan scoring, Katrien De Bock and Ewald lab members for critical discussions of the manuscript, Ann Rougvie for pJA1 [P*col-19*::GFP] plasmid, Michel Labouesse for OD76, ML2600 strains, Alex Hajnal for AH4617, AH3437, AH3284 strains, Cathy Savage-Dunn for CS637, CS678, Harald Hutter for VH2847 strain, Taina Pihlajaniemi for TU1 strain, WormBase for curated gene and phenotype information, Tobias Schwarz and the support of the Scientific Center for Optical and Electron Microscopy ScopeM of the Swiss Federal Institute of Technology ETHZ. Some strains were provided by the CGC, which is funded by the NIH Office of Research Infrastructure Programs (P40 OD010440). Figure 3N was created with BioRender.com (Publication license JH23SSQB6M). Funding from the Swiss National Science Foundation PP00P3_163898 and 190072 to ACT, CS, AG, and CYE, and the European Research Council AdvG grant 670821 to RA is acknowledged.

---

**Supplementary Figure 1.**
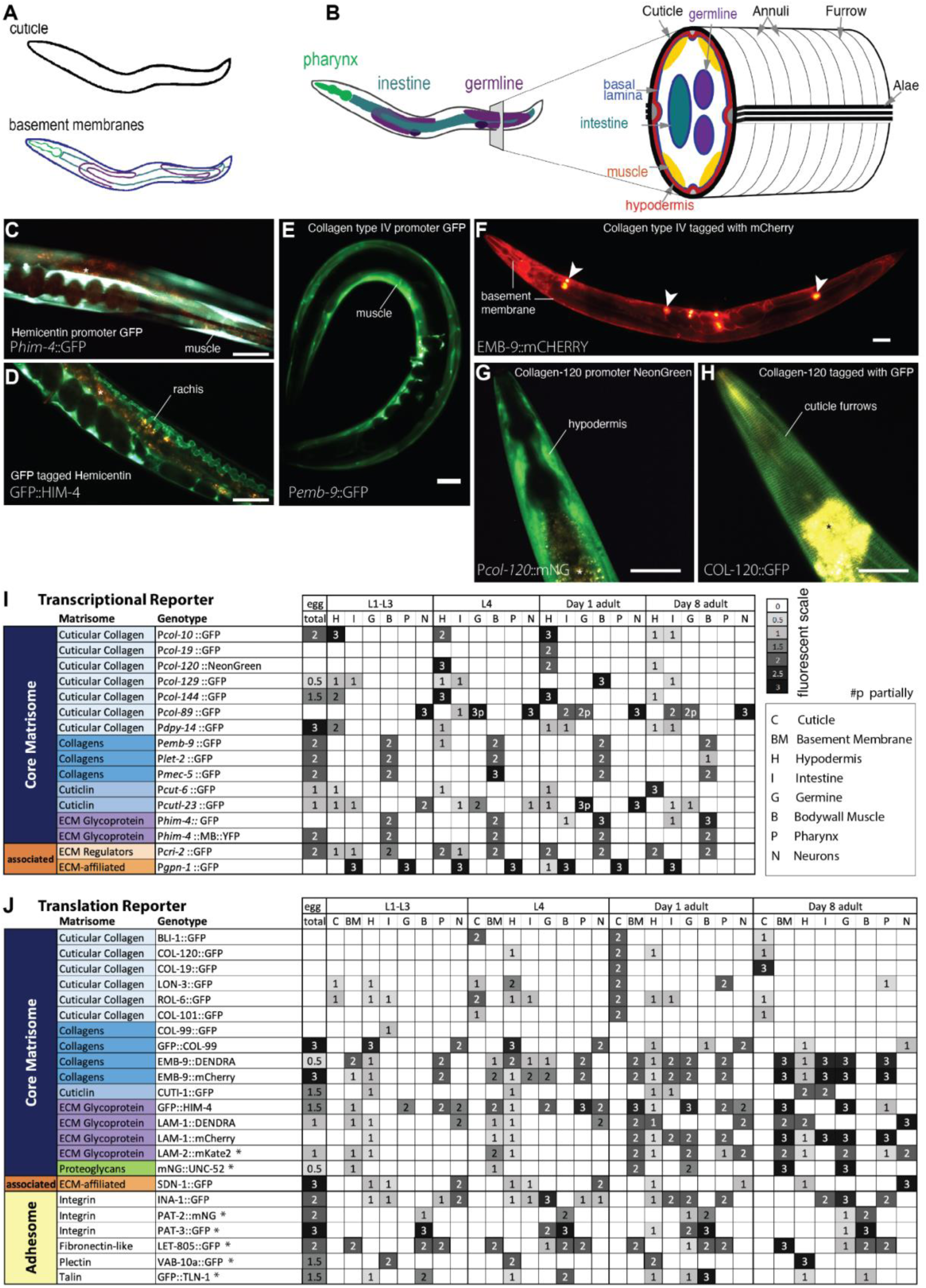
Age-associated changes of matrisome proteins in the ECM. (A, B) Schematic of *C. elegans’* tissues and ECMs (basement membrane and cuticle). (C, D) Expression of hemicentin *him-4* promoter-driven in body wall muscles (C) and GFP tagged HIM-4 protein incorporated in ECM (D). (E, F) Expression of collagen type IV *emb-9* promoter-driven in body wall muscles (E) and mCherry tagged EMB-9 protein incorporated in almost all basement membranes surrounding pharynx, intestine, gonad, and apical surface of body wall muscles (F). Arrowheads indicate coelomocytes, the *C. elegans’* macrophage-like cells, which scavenge foreign particles including fluorescent proteins from the pseudocoelomic fluid. (G, H) Expression of cuticular collagen *col-120* promoter-driven in the hypodermis (G) and GFP tagged COL-120 protein incorporated in the cuticular furrows (H). (I) Transcriptional reporters are driven by matrisome genes during development and aging. For details, see Supplementary Table 1. The fluorescent scale corresponds to the highest observed fluorescence of a reporter line (intensity 3) graded to no observed fluorescence above background (intensity 0). (J) Translational reporters of matrisome and adhesome proteins are localized and incorporated into ECM structures during development and aging. For details, see Supplementary Table 1. * indicates CRISPR-Cas9 genome inserted tag in the endogenous gene locus. (C-H) scale bar = 50 µm, (C,D,G,H) * autofluorescent gut granules are in brown-yellowish.

**Supplementary Figure 2.**
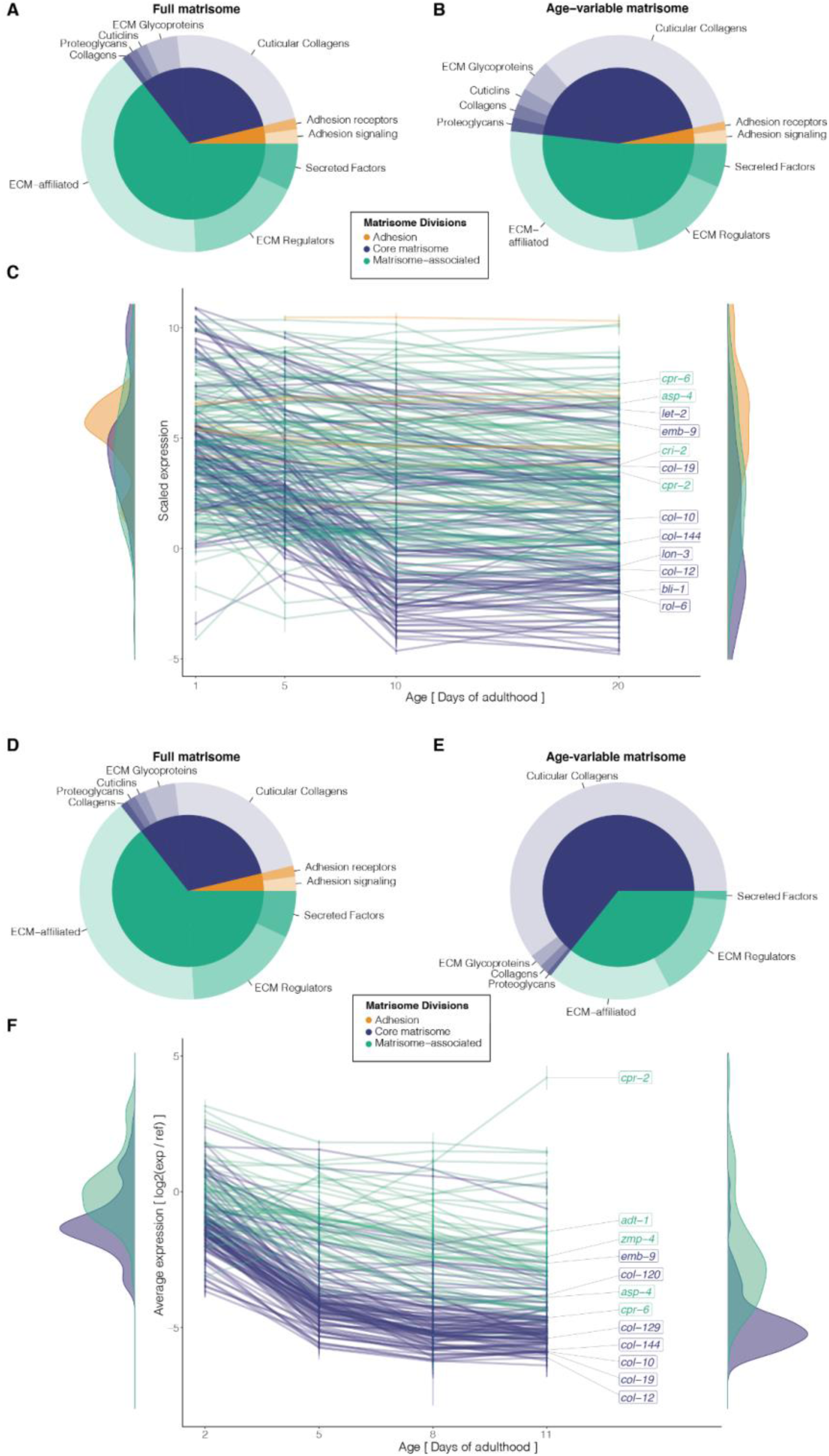
Transcriptional changes of matrisome and adhesome during aging. (A, B) Composition of the matrisome and adhesome (A) and their age-variable subset (B) based on the GSE12168 expression profile (B). (C) Longitudinal expression of matrisome and adhesome genes during aging (GSE12168). (D, E) Composition of the matrisome and adhesome (D) and their age-variable subset (E) based on the GSE46051 expression profile (E). (F) Longitudinal expression of matrisome and adhesome genes during aging (GSE46051). (A-F) For details, see Supplementary Table 2.

**Supplementary Figure 3.**
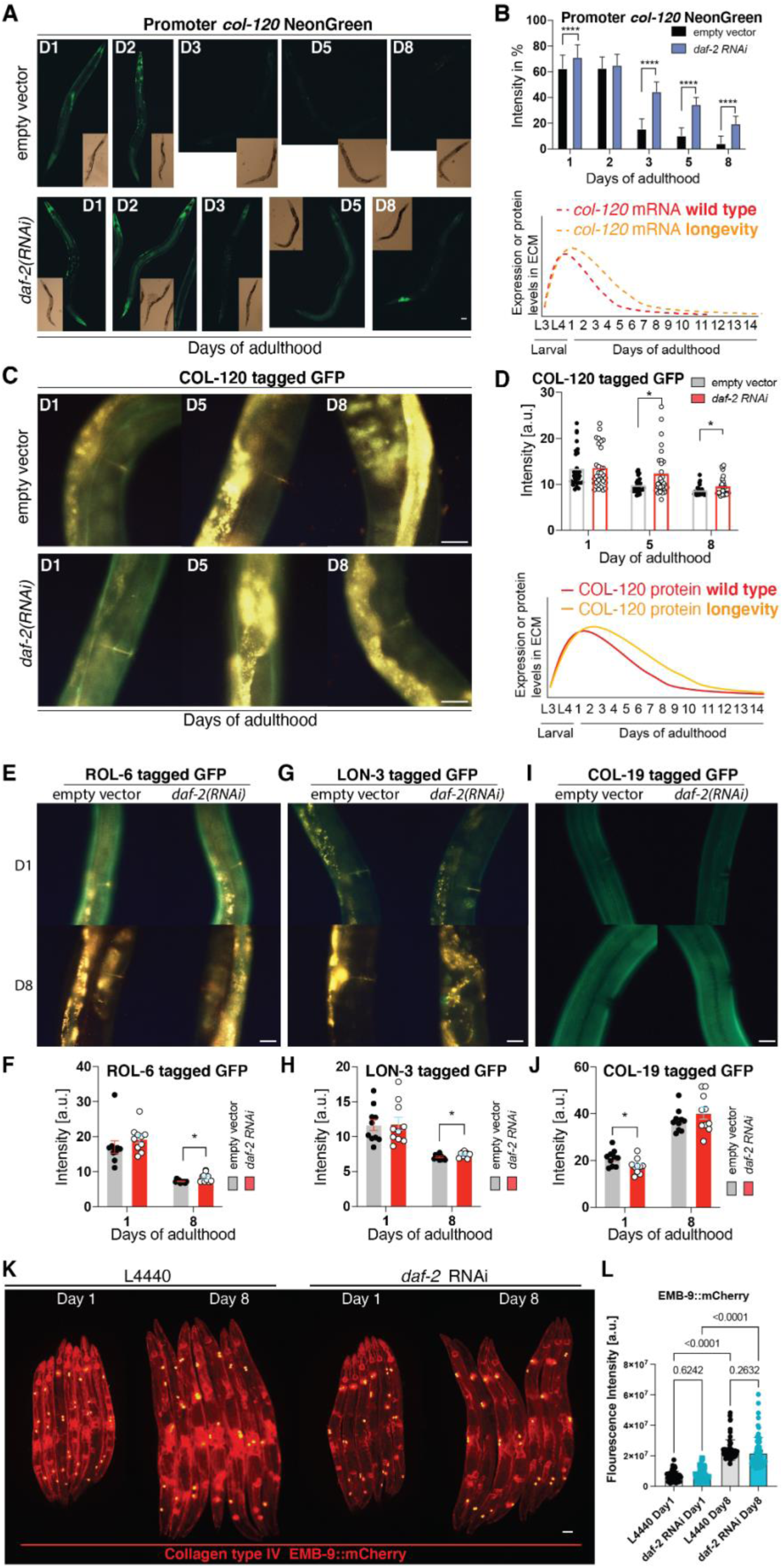
Collagen levels in the ECM during aging and longevity. (A) Time course of LSD1107 *Pcol-120*::NeonGreen animals fed with *daf-2* RNAi or the empty RNAi vector control L4440 bacteria. Scale bar = 50 µm. (B) Quantification and model of the LSD1107 P*col-120*::NeonGreen time course. Three rounds, each n=20, were quantified using a visual grading scale system with values from 0 - 3 in 0.5 steps. Error bars represent SDs. For details and data, see Supplementary Table 4. (C) Images of LSD2043 COL-120::GFP animals fed with *daf-2* RNAi or the empty RNAi vector control L4440 bacteria during aging, taken on day 1, day 5, and day 8 of adulthood. Scale bar = 25 µm (D) Quantification of LSD2043 COL-120::GFP green fluorescence intensity levels shown as a composite of 3 independent biological trials of each about 10 *C. elegans* per condition and day. For details and data, see Supplementary Table 4. Below is the model of COL-120 levels in ECM during aging and upon longevity. (E, G, I) Images of LSD2022 ROL-6::GFP, LSD2063 LON-3::GFP, and LSD2064 COL-19::GFP animals fed with *daf-2* RNAi or the empty RNAi vector control L4440 bacteria during aging, taken on day 1 and day 8 of adulthood. Scale bar = 25 µm. (F, H, J) Green fluorescence intensity quantification of two separate experiments, each three rounds of *daf-2* or L4440 RNAi experiments on translational cuticular collagen reporter strains. The green fluorescence intensities (excluding autofluorescence) of images of either 3 independent trials of about 10 animals (LON-3, COL-19) or 1 trial of 10 (ROL-6) animals were quantified (see Materials and Methods for details). The data are represented as mean and SD. * indicates *P*-value ≤ 0.05 determined by using a two-way ANOVA. For details and data, see Supplementary Table 4. (K- L) An orthologue of mammalian Type IV collagen EMB-9 increases with age independent of slowing aging upon reduced Insulin/IGF-1 signaling. (K) Representative images of EMB::mCherry animals treated from eggs with empty vector (L4440) or *daf- 2*(RNAi) and scored at day 1 and day 8 of adulthood at 20°C. Scale bar = 50 µm. (L) Quantification of EMB-9::mCherry fluorescent intensity. Each dot represents an animal. 3 independent biological trials. *P*-value determined with One-way ANOVA. For raw data and statistics, see Supplementary Table 4.

**Supplementary Figure 4.**
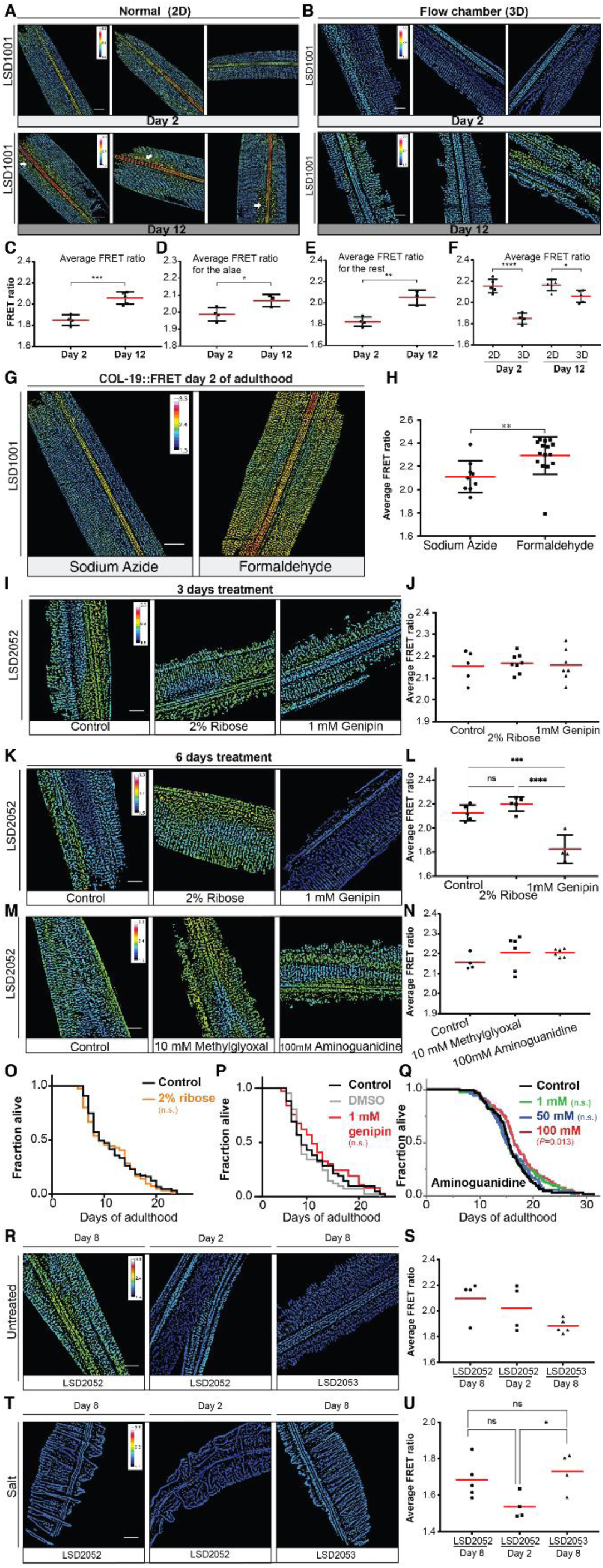
Collagen FRET-reporter implicates tissue tension and crosslinking associated with aging. (A-F) Comparison of FRET ratio images of transgenic LSD1001 COL-19::FRET animals imaged using normal (2D) or our developed flow chambers (3D) on day 2 and day 12 of adulthood and corresponding quantification. (G) Representative FRET ratio images of transgenic LSD1001 COL-19::FRET animals anesthetized with 25 mM sodium azide or fixed with 4% formaldehyde. Scale bar = 20 μm, FRET calibration bar between 1.5 - 3.3, (H) Quantitative analysis of FRET ratios from the whole cuticle revealed significantly higher FRET ratios for LSD1001 COL-19::FRET *C. elegans* fixed with 4% formaldehyde for 24 hours compared to LSD1001 COL::FRET *C.elegans* than were anesthetized with 25 mM sodium azide. (I-N) Representative FRET ratio images and quantification of transgenic LSD2052 COL- 19::FRET animals treated from day 1 of adulthood with different chemicals and scored at day 4 (I, J) or day 7 of adulthood (K-N). (O-Q) The lifespans of *C. elegans* were treated with chemicals starting during adulthood (Raw data and statistical details in Supplementary Table 7). (R-U) Lowering inner pressure by osmotic conditions. On day 2 or 8 of adulthood, either normal-lived (LSD2052) or longevity-promoting (LSD2053) transgenic COL-19::FRET animals were picked and placed directly either into a physiological buffer or high sodium chloride (1g NaCl/ 10mL M9 buffer) containing flow chambers and then imaged. (C-F, H, J, L, N, S, U) Error bars correspond to the standard deviation of the mean, * *P* < 0.05, ** *P* < 0.01, *** *P* < 0.001, **** *P* < 0.0001. Statistically significant differences between mean values were calculated using an unpaired *t*-test. For raw data, details, and statistics, see Supplementary Table 6.

**Supplementary Figure 5.**
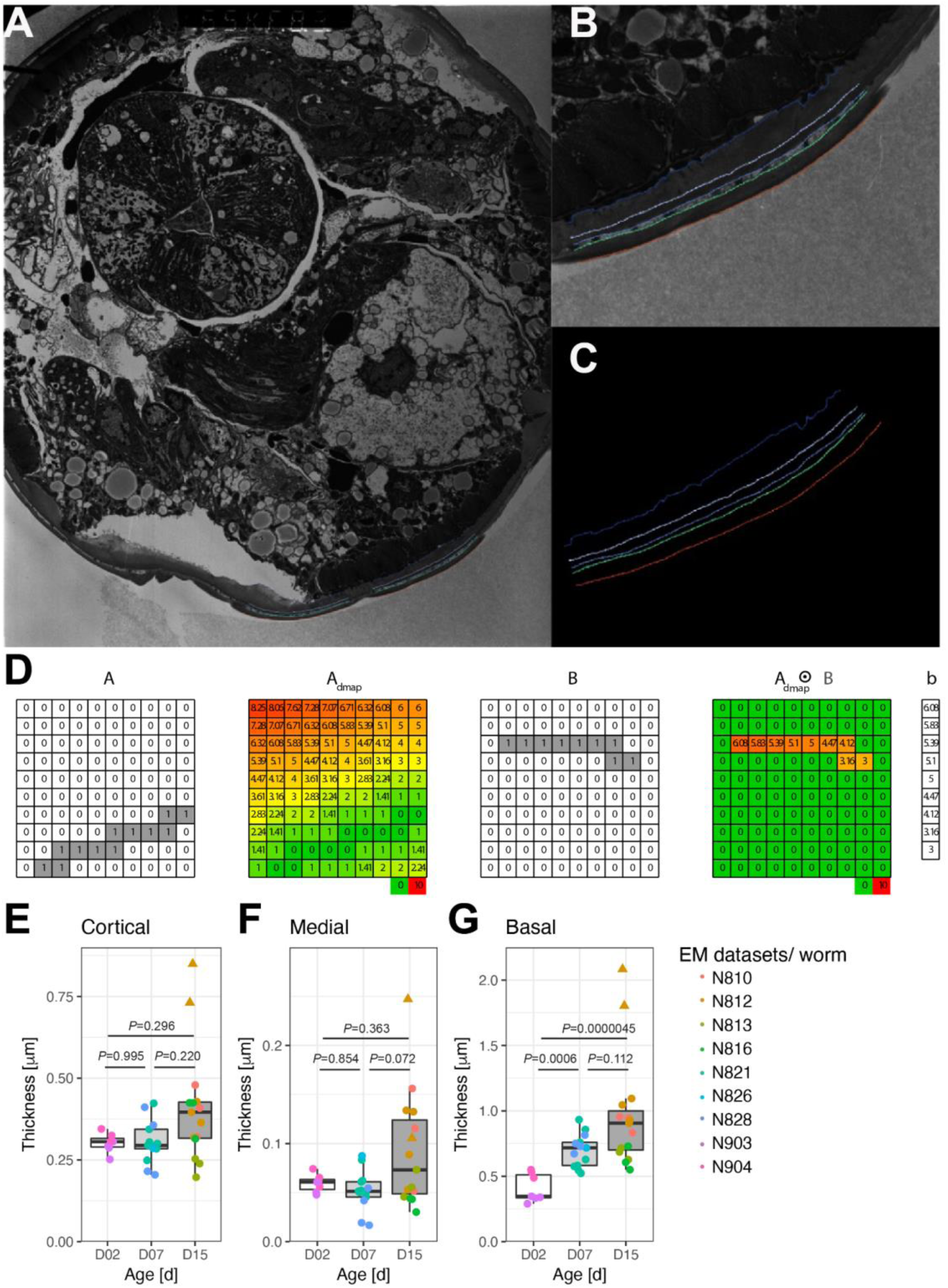
Cuticle thickness increases during aging. (A) Transverse transmission electron micrograph of a 15 days old adult (Source: wormimage.org). Colored curves were drawn manually in regions included in the analysis of cuticle thickness. (B) Zoom into a region used for evaluation. (C) Colored curves correspond to binary masks exported to Matlab for evaluation. (D) Workflow to measure the thickness of annotated cuticle layers. . A = Binary mask of first layer border. Admap = Distance transform of A. B = Binary mask second layer border. Admap ⊙ B = Elementwise multiplication of Admap and B, non-zero values correspond to the shortest distances to A for every pixel in B. (E-G) Quantification of the thickness of (E) cortical (F) medial and G) basal cuticle. Dots correspond to individual animals. Triangles are the outliers. Boxplot shows the median (black line), 25th/75th percentiles (hinges), and 1.5*IQR (whiskers). *P*-values are One- way ANOVA post hoc Tuckey without including outliers. See Figure 3E for total cuticle thickness and Supplementary Table 9 for details.

**Supplementary Figure 6.**
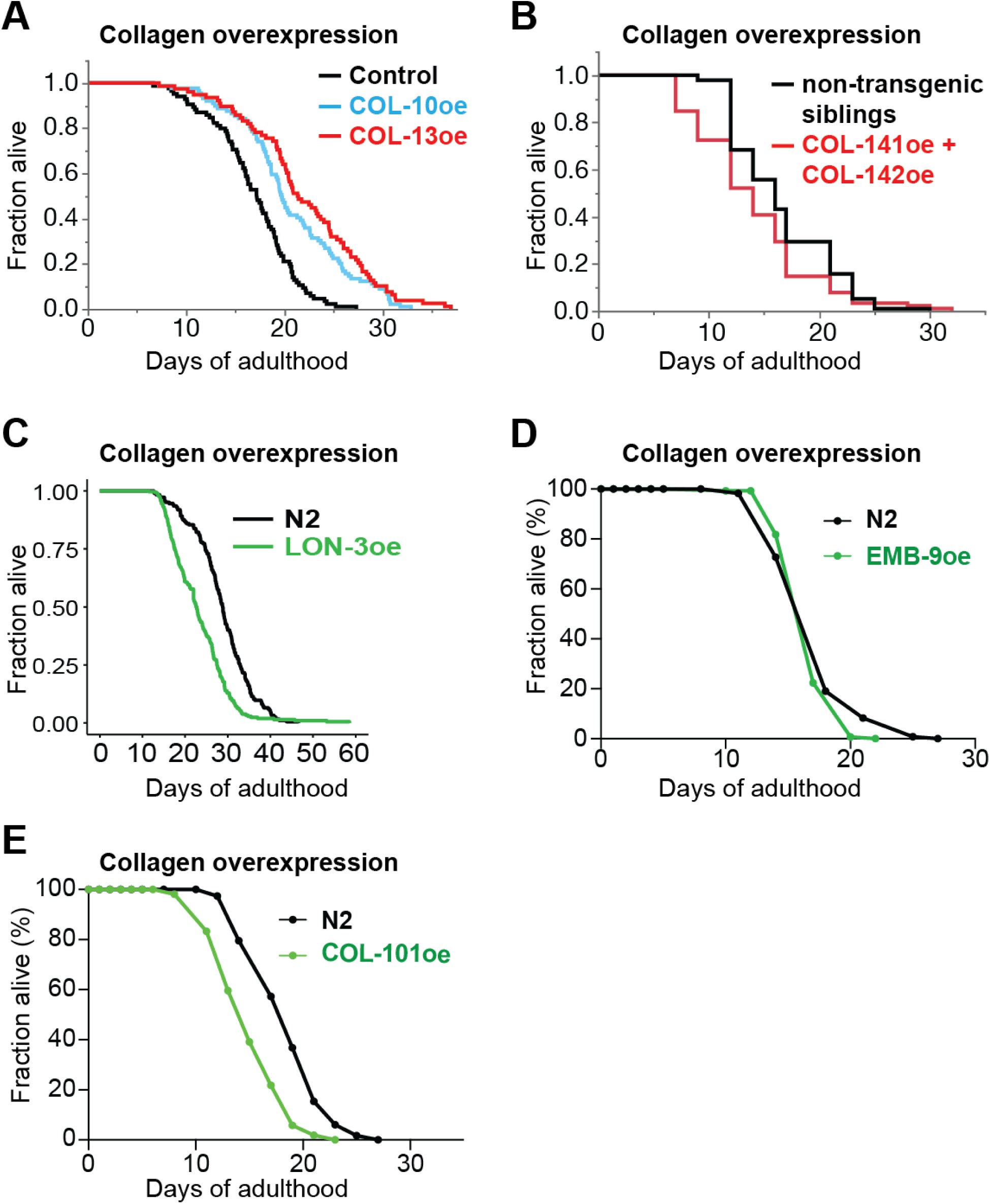
Only key collagen overexpression is sufficient to increase the lifespan. (A) Cuticular collagens COL-10 (LSD2018) or COL-13 (LSD2014) overexpression (oe) increased lifespan compared to control (wild type with *rol-6(su1006)* co-injection marker LSD2013) on UV-inactivated bacteria. (B) Cuticular collagens COL-141+COL-142 (CS637) overexpression (oe) did not extend lifespan compared to non-transgenic sibling control. (C) Cuticular collagen LON-3 (*kuIs55*) overexpression (oe; 8x outcrossed to N2) did not extend lifespan compared to wild type (N2). (D) Basement membrane Type IV collagen EMB-9 (NK364) overexpression (oe) did not increase lifespan compared to wild type (N2). (E) Cuticular collagen COL-101 (*dmals40*) overexpression (oe) shortened lifespan compared to wild type (N2). (A-E) For details, raw data, and statistics, see Supplementary Table 7.

**Supplementary Figure 7.**
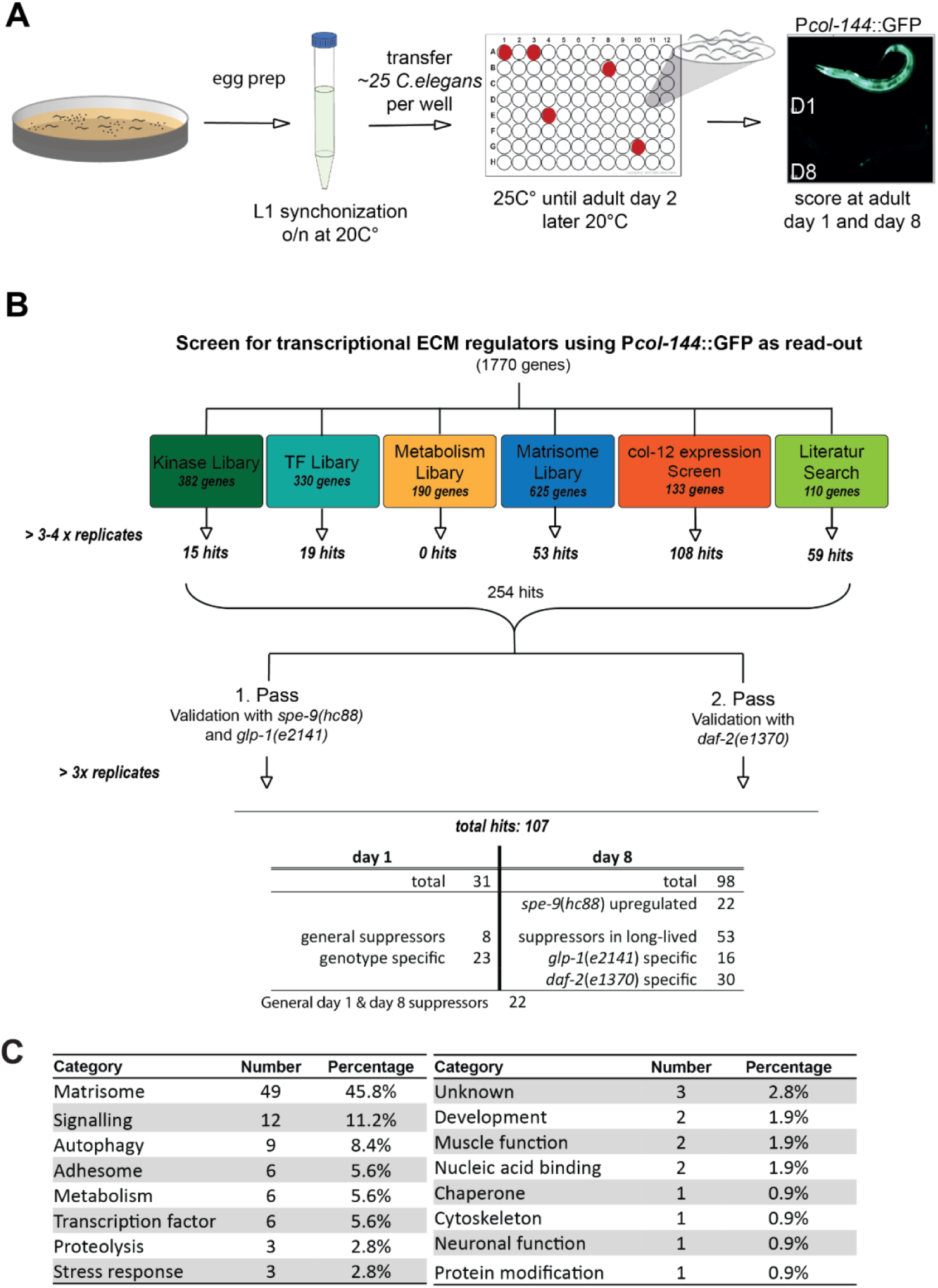
RNAi screen set up and results. (A) Schematic workflow of the screen. (B) Schematic overview of the targeted RNAi screen. 1770 RNAi clones were screened in a first pass using transgenic *C. elegans* strains expressing the P*col-144*::GFP reporter in either a *spe-9*(*hc88*) or a long-lived *glp-1*(*e2141*) background. These hits were re- validated in *spe-9*(*hc88*) and *glp-1*(*e2141*) and additionally in *daf-2*(*e1370*) animals identifying 107 high confident hits. (C) Table of confident hits of RNAi screen for transcriptional ECM regulators sorted by gene ontology categories, number of hits per category, and the percentage of each category being represented in relation to all screen hits. For the categorization, the WormCat online tool was used. For details, see Supplementary Table 10.

**Supplementary Figure 8.**
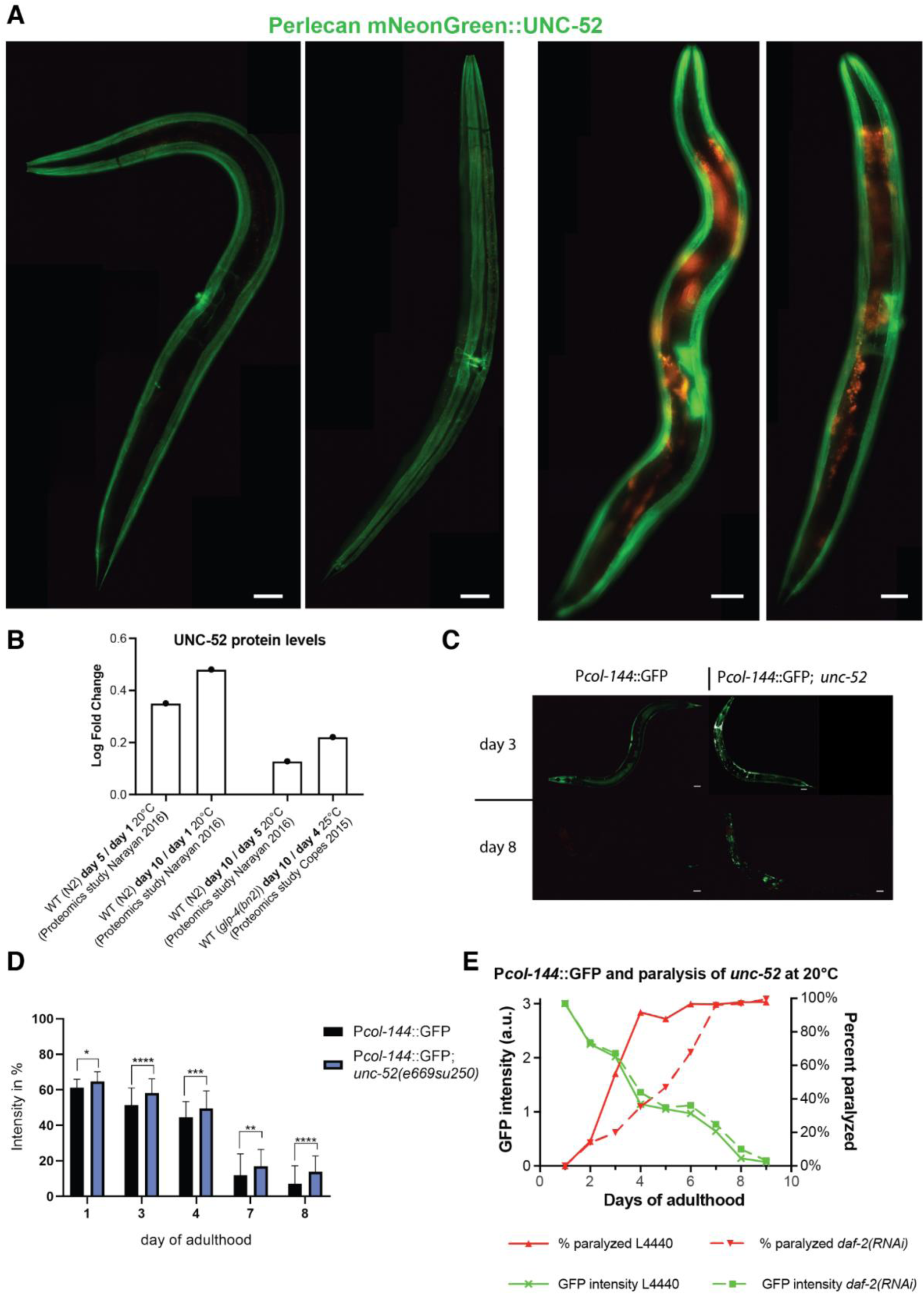
Perlecan levels during aging and functional consequences of loss of perlecan integrity. (A) Fluorescent microscopy images of CRISPR inserted NeonGreen tagging perlecan (NK2583 *unc-52*(*qy80* [NeonGreen::UNC-52]) at days 1 and 8 of adulthood. Scale bar = 50 μm. (B) Relative increase of UNC-52 perlecan levels during aging quantified from previous proteomics studies (See Supplementary Table 4 for details). (C) Fluorescent images of P*col-144*::GFP; *unc-52(e669su250)* mutants at day 3 and 8 of adulthood at 20°C compared with P*col-144*::GFP in wild-type background. Scale bar = 50 µm. (D) Quantification of P*col-144*::GFP fluorescence in *unc-52(e669su250)* temperature-sensitive mutants at semi-permissive 20°C during aging. Three rounds, each n=20. See Supplementary Table 12 for statistics and details. (E) Longevity intervention *daf-2(RNAi)* postponed the perlecan *unc-52(e669, su250)* mutant paralysis phenotype (dashed red line) but failed to prolong collagen expression (P*col-144*::GFP, green dashed line) during aging at semi-permissive temperature 20°C. For details, see Supplementary Table 12.

**Supplementary Figure 9.**
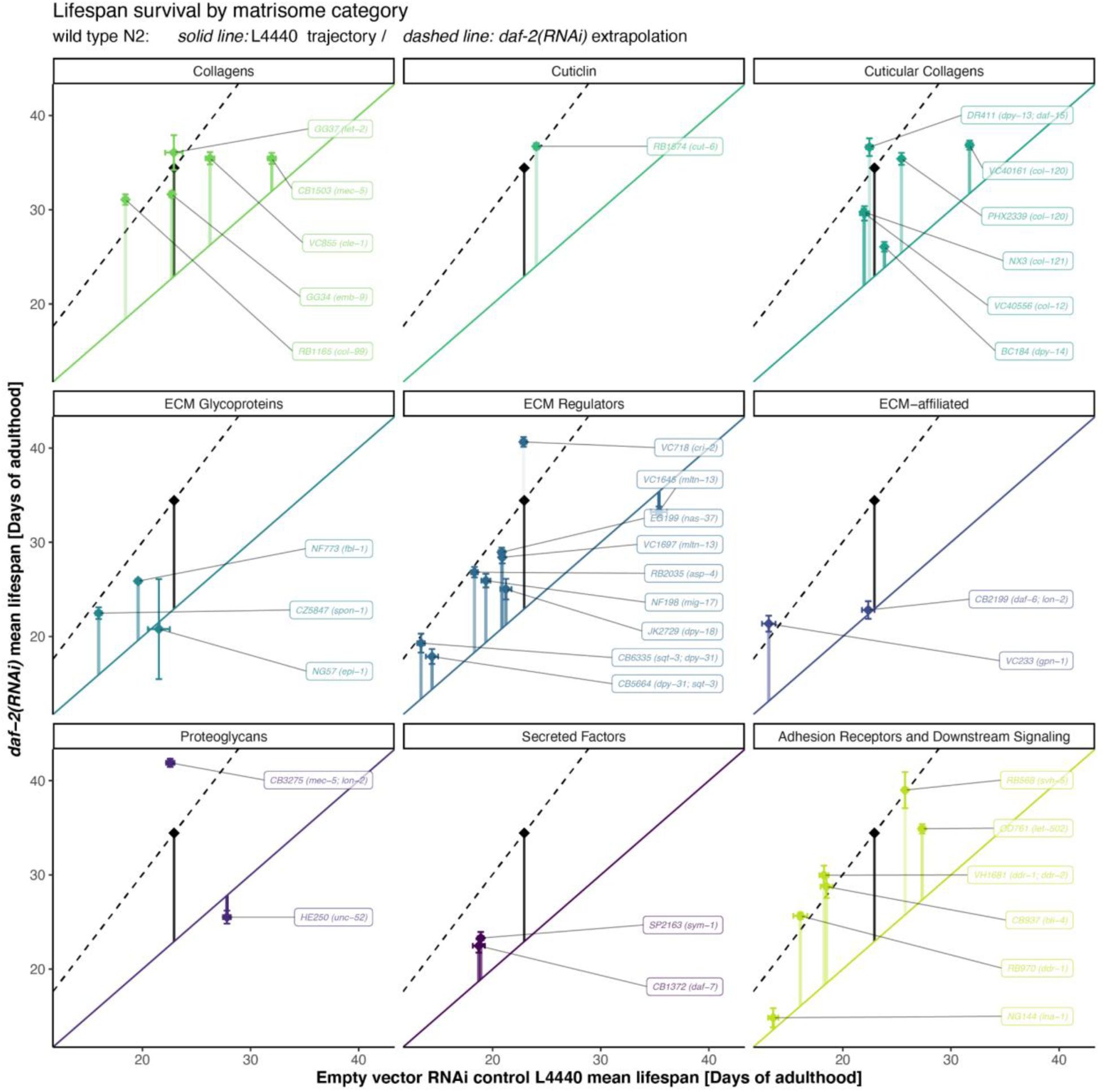
All matrisome categories are implicated in promoting healthy aging. Lifespan differences between EV (x-axis) and *daf-2 (y-axis)* RNAi treatment are displayed for all matrisome mutants grouped by matrisome category. The solid diagonal (y = x) represents the ‘no effect line’ and the dashed line represents the lifespan ratio of the overall global wild type across all runs comparing *daf-2* vs. EV RNAi. Each genotype is labeled in its corresponding facet and color while the wild type is shown in black. All details, raw data, and statistics are in Supplementary Table 7.

**Supplementary Figure 10.**
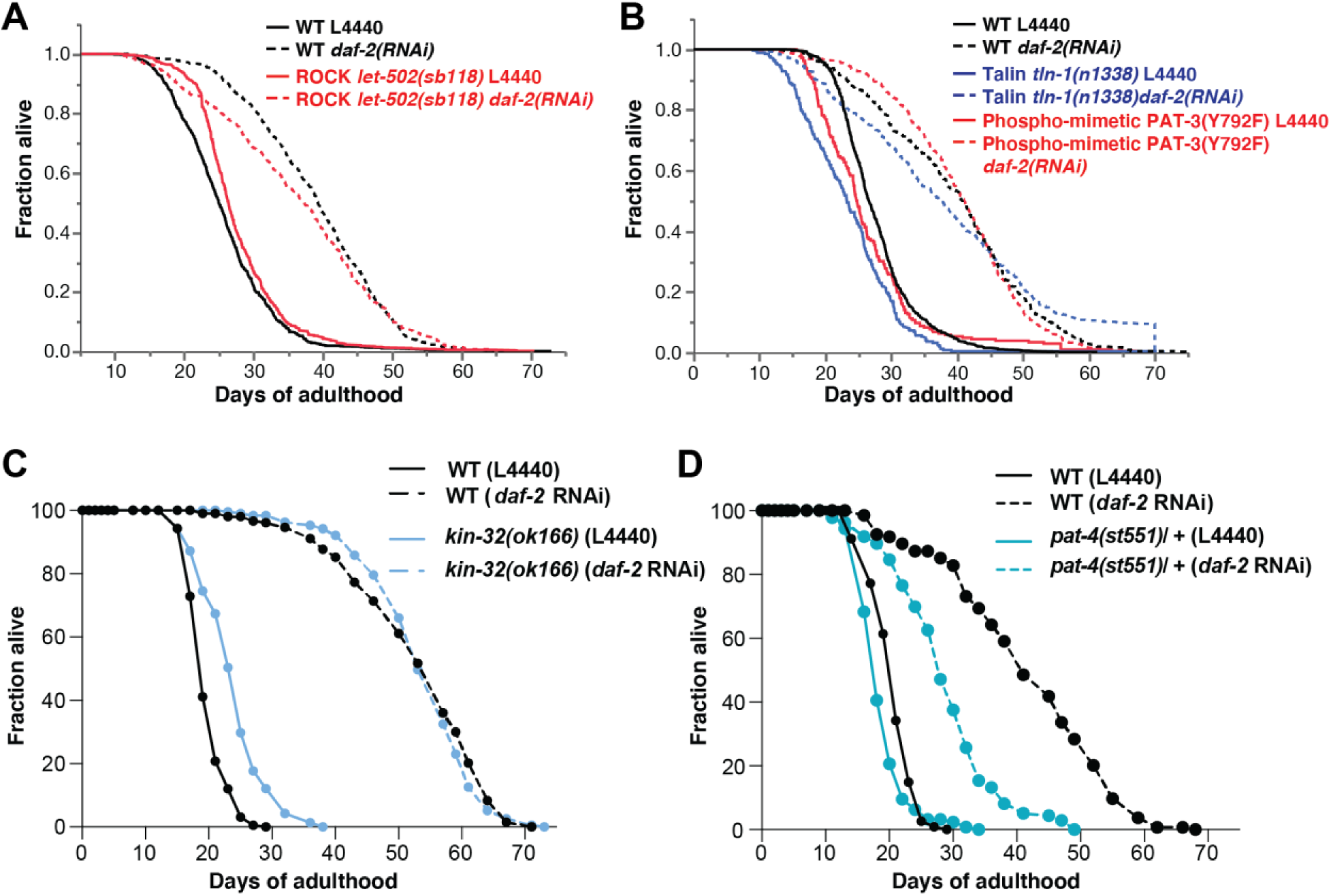
Mechanotransductive signaling genes and their functional role in longevity. (A) The integrin downstream ROCK kinase was not required for longevity at 20°C. (B) Both the phospho-mimetic PAT-3(Y792F) that activates talin signaling and *tln- 1(n1338)* mutants were not required for longevity at 20°C. (C) Mutation in focal adhesion kinase *kin-32(ok166)* increased lifespan compared to wild type but did not suppress *daf-2(RNAi)* mediated longevity at 20°C. (D) Heterozygous lethal mutation in *pat-4(st551)*/+ shortened WT lifespan and longevity upon reduced Insulin/IGF-1 receptor signaling at 20°C. (A-D) For raw data and statistical details, see Supplementary Table 7.

## Materials and Methods

### C. elegans strains

*Caenorhabditis elegans* strains were grown on NGM plates with OP50 *Escherichia coli* bacteria at 20°C as descripted in (Stiernagle, 2006). The Bristol N2 was used as a wild-type *C. elegans* strain (Brenner, 1974). Most strains were obtained from the Caenorhabditis Genetics Center [CGC]: BC184 *dpy-14(e188) unc-13(e51) bli-4(s90)/unc- 15(e73)* I, BC10074 *dpy-5*(*e907*) I; *sEx10074*[P*emb-9*::GFP + pCeh361], BC11902 *dpy-5*(*e907*) I; *sEx11902* [P*col-129*::GFP + pCeh361], BC12229 *dpy-5*(*e907*) I; *sEx10002* [P*cutl-23*::GFP + pCeh361], BC12275 *dpy-5*(*e907*) I; *sEx12275* [P*cut-6*::GFP::GFP + pCeh361], BC12533 *dpy-5*(*e907*) I; *sEx12533* [P*col-89*::GFP + pCeh361], BC12900 *dpy- 5*(*e907*) I; *sIs11600* [P*mec-5*::GFP+ pCeh361], BC13149 *dpy-5*(*e907*) I; *sEx13149* [P*him-4*::GFP + pCeh361], BC13560 *dpy-5*(*e907*) I; *sIs13559* [P*col-59*::GFP + pCeh361], BC13623 *dpy-5*(*e907*) I; *sEx13623* [P*cri-2*::GFP + pCeh361], BC13861 *dpy-5*(*e907*) I; *sIs13252* [P*let-2*::GFP + pCeh361], BC14295 *dpy-5*(*e907*) I; *sEx14295* [P*gpn-1*::GFP+ pCeh361], CB937 *bli-4(e937)* I, OD761 *let-502(sb118)* I, MT3100 *tln-1(n1338)* I, CB698 *vab-10(e698)* I, VC117 *vab-10(gk45)* I, GOU2043 *vab-10a(cas602*[*vab-10a*::gfp]) I, RB776 *kin-32(ok166)* I, VC855 *cle-1(gk364)* I, CZ5847 *spon-1(ju402)* II; juEx1111, HE250 *unc-52(e669su250)* II, BT24 *rhIs23* [GFP::HIM-4] III, CB1372 *daf-7(e1372)* III, GG37 *emb-9(g34)* III, JK2729 *dpy-18(ok162)* III, RB1574 *cut-6(ok1919)* III, RW1522 *pat-2(st538) unc-32(e189)* III; *stEx10* [*pat-2(+); rol-6(su1006)*], RW3550 *pat-4(st551)/unc- 45(e286)* III, NJ268 *pat-3(rh96)* III, NG144 *ina-1(gm144)* III, HE250 *unc-52(e669, su250)* III, CH1445 *unc-119*(*ed3*) III; *cgEx198* [BLI-1::GFP + *unc-119*(*+*)], HS428 *dpy-22*(*os26*) X; *osEx89* [COL-10::GFP + *dpy-22*(*+*)], MH2051 *kuIs55* [LON-3::GFP + *unc-119*(*+*)]; pYSL3G3, NG2517 *him-5*(*e1490*) V; [INA-1::GFP + *rol-6*(*su1006*)], NK248 *unc-119*(*ed4*) III; *qyIs10* [LAM-1::GFP + *unc-119*(*+*)] IV, NK2583 *unc-52*(*qy80* [NeonGreen::UNC-52]), NK358 *unc-119*(*ed4*) III; *qyIs43* [PAT-3::GFP + INA-1(genomic) + *unc-119*(*+*)], NK364 *unc-119*(*ed4*) III; *qyIs46* [EMB-9::mCherry + *unc-119*(*+*)], NK651 *unc-119*(*ed4*) III; *qyIs108* [LAM-1::Dendra + *unc-119*(*+*)], NK696 *unc-119*(*ed4*) III; *qyIs127* [LAM-1::mCherry + *unc-119*(*+*)], NK860 *unc-119*(*ed4*) III; *qyIs161* [EMB-9::Dendra + *unc- 119*(*+*)], ML2501 *let-805(mc73*[*let-805*::gfp + *unc-119*(+)]) *unc-119(ed3)* III, NK2446 *qy41* [*lam-2*::mKate2] X, NK2479 *qy49*[*pat-2*::2xmNG] III, RP247 *trIs30* [P*him-4*::MB::YFP + P*hmr-1b*::DsRed2 + P*unc-129nsp*::DsRed2], TP12 *kaIs12* [COL-19::GFP], WS3403 *opIs170* [SDN-1::GFP::*unc-54* 3’UTR + *lin-15*(*+*)], WT30 *unc-119*(*ed3*) III; *wtEx30* [CUTI- 1::GFP + *unc-119*(*+*)], TJ1060 *spe-9(hc88)* I; *rrf-3(b26)* II, CB4037 *glp-1(e2141)* III, LD1036 *daf-2(e1370); him-8(e1489),* DR411 *dpy-13(e184)/daf-15(m81)* IV, NF773 *fbl- 1(k201)* IV, NG57 *epi-1(gm57)* IV, NX3 *col-121* IV, RB1165 *col-99(ok1204)* IV, TM8818 *col-101(tm8818)* IV, VC40161 *col-120* IV, CB5664 *dpy-31(e2770)* III; *sqt-3(e2809)* V, CB6335 *dpy-31(e2919)* III; *sqt-3(e2906)* V, NF198 *mig-17(k174)* V, TM6673 *col- 10(tm6673)* V, VC718 *cri-2(gk314)* V, VC40556 *col-12* V, BT12 *him-4(rh319)* X, CB1503 *mec-5(e1503)* X, CB2199 *lon-2(e678); daf-6(e1377)* X, CB3275 *lon-2(e678) mec-5(e1504)* X, CH1878 *dgn-2(ok209) dgn-3(tm1092) dgn-1(cg121)* X; *cgEx308* [pJK600/dgn-1(+) + pJK602/*dng-1*p::GFP + *rol-6(su1066)*], EG199 *nas-37(ox199)* X, GG37 *let-2(g37)* X, RB970 *ddr-1(ok874)* X, RB2035 *asp-4(ok2693)* X, RB568 *svh- 5(ok286)* X, RT3574 *lin-15B&lin-15A(n765)* X; *ihIs35* [YAP-1::GFP::*unc-54* 3’UTR + *lin-15*(+)], SP2163 *sym-1(mn601)* X, VC233 *gpn-1(ok377)* X, VC1645 *mltn-13(gk766)* X, VC1697 *mltn-13(gk807)* X, *yap-1(tm1416)* X.

The strains AH3284 *pat-3(zh105[PAT-3(Y792F)])*, AH3437 *zh117* [GFP::TLN-1], AH4617 *zh115* [PAT-3::GFP] III (Walser et al., 2017), DMS1020 *dmaIs40* [*col-101*p::*col-101*::GFP (40 ng/μl); *unc-54*p::mCherry (40 ng/μl)] (Zhang et al., 2020), OD761 *let-502(sb118)* (Diogon et al., 2007), ML2600 *vab-10*(*mc100*[VAB-10A::mCherry+loxp])I (Suman et al., 2019), CS678 *col-141(lf)*, CS637 Ex [COL-141COL-142(oe); Pmyo-2::GFP], *jgIs5* [ROL-6::GFP;TTX-3::GFP] (Kim et al., 2011), TU1 *unc-119*(*ed3*)III; [P*col-99*[16655]::S0001_pR6K_Amp_2xTY1ce_EGFP_FRT_rpsl_neo_FRT_3xFlag] dFRT::unc-119 (Tu et al., 2015), LSD1001 [*pha-1(e2123)*; *col-19*(FRET between exon)- version C3 + PHA-1(+)] (Meng et al., 2008), VH2847 *hdIs73* [COL-99::GFP, pha-1(+)] (Taylor et al., 2018) were gifts from other labs.

The last group of strain was generated by UV integration, crossing, or injection in our lab: LSD1000 *xchEx001* [(pRedFlp-Hgr)(*col- 120*[30044]::S0001_pR6K_Amp_2xTY1ce_EGFP_FRT_rpsl_neo_FRT_3xFlag)dFRT::u nc-119-Nat]; pRF4 [*rol-6(su1006)*], LSD1106 *pha-1*(*e2123*) III; *xchEx105* [P*col- 120*::NeonGreen; *pha-1*(*+*)], LSD1107 xchEx017 [P*col-120*::NeonGreen; P*col- 12*::DsRed], LSD2001 *xchIs001* [P*col-144*:: GFP; *pha-1*(+)], LSD2043 *xchIs012* [(pRedFlp-Hgr) (*col-120* [30044]::S0001 pR6K Amp 2xTY1ce EGFP FRT rpsl neo FRT 3xFlag) dFRT::unc-119-Nat]; pRF4 *rol-6(su1006gf)*, LSD2051 [*col-19*(FRET between exon)-version C3 + PHA-1(+)] was made by UV integration of LSD1001 and outcrossing 12 times, LSD2052 *spe-9(hc88)*; [*col-19*(FRET between exon)-version C3 + PHA-1(+)], LSD2052 *pha-1(e2123)*; [col-19(FRET between exon)-version C3]; *spe-9(hc88)* I, LSD2053 *glp-1(e2141)*; [*col-19*(FRET between exon)-version C3 + PHA-1(+)], LSD2063 *kuIs55* [LON-3::GFP + *unc-119*(+) pYSL3G3 rollers]; *spe-9(hc88)* I, LSD2117 *xchIs016* [P*col-19*::GFP], LSD2002 *spe-9(hc88)* I ; *xchIs001* [P*col-144*:: GFP; *pha-1(+)*], LSD2003 *glp-1*(*e2141*) III ; *xchIs001* [P*col-144*:: GFP; *pha-1*(*+*)], LSD2117 *xchIs016* [P*col- 19*::GFP], LSD2122 *spe-9(hc88)* I; *daf-2(e1370)* III; *xchIs001* [P*col-144*:: GFP; *pha-1(+)*], LSD1061 *xchEx062* [P*col-120*::*col-120*::dendra2; pha-1 (+)]; pha-1(e2123)III, LSD2191 *xchIs001* [P*col-144*:: GFP; *pha-1(+)]* X; *unc-52(e669su250)*, LSD2197 *xchIs001* [P*col- 144*:: GFP; *pha-1(+)*] X; *pat-3(rh96) III*, LSD2147 *qyIs46* [P*emb-9*::*emb-9*::mCherry + *unc- 119(+)]* ; *pat-3(zh115* [PAT-3::GFP]); unc-119(ed4) III, LSD2161 *qyIs46* [P*emb-9*::*emb-9*::mCherry + *unc-119(+)]; unc-52(e669su250) II* ; *pat-3(zh115* [PAT-3::GFP]); *unc- 119(ed4)* III, LSD2207 *vab-10*(*mc100*[VAB-10A::mCherry+loxp])I; *ihIs35* [YAP-1::GFP::*unc-54* 3’UTR + *lin-15*(+)].

### Cloning of transgenic constructs

The plasmid pXCH8 (P*col-120*::NeonGreen) was built and purchased from Vectorbuilder. The promoter sequence originates from *col-120* fosmid clone WRM0622A_D10(pRedFlp- Hgr)(col- 120[30044]::S0001_pR6K_Amp_2xTY1ce_EGFP_FRT_rpsl_neo_FRT_3xFlag)dFRT::u nc-119-Nat from the TransgeneOme Project (Sarov et al., 2012), while the plasmid DG398 (Slot2 ENTRY vector for mNeonGreen::3xFlag) (Hostettler et al., 2017) served as backbone.

### Generation of mutant *C. elegans* strains

PHX2339 *col-120(syb2339)* was generated by SunyBiotech, and is a 1085 deletion of the Y11D7A.11 covering the ATG to the TAA stop and was 4 times outcrossed.

### Generation of transgenic lines

LSD1106 *pha-1*(*e2123*) III; *xchEx105* [P*col-120*::NeonGreen; *pha-1*(*+*)]. Generated by injecting 50 ng/μl pXCH8 (P*col-120*::NeonGreen) with 50 ng/μl pBX (*pha-1* (+)) co- injection marker into *pha-1*(*e2123*)III mutants. LSD1107 *xchEx017* [P*col- 120*::NeonGreen; P*col-12*::DsRed] was generated by injecting 50 ng/μl pXCH8 (P*col- 120*::NeonGreen) with 50 ng/μl P*col-12*::DsRed into N2.

LSD2001 *xchIs001* [P*col-144*:: GFP; *pha-1*(+)] was generated by integration of Ex [P*col- 144*:: GFP; pha-1(+)] (Budovskaya et al., 2008) via UV light irradiation and 8 times outcrossing with N2 animals.

The strain LSD2043 *xchIs012* [(pRedFlp-Hgr) (col-120 [30044]::S0001 pR6K Amp 2xTY1ce EGFP FRT rpsl neo FRT 3xFlag) dFRT::unc-119-Nat]; pRF4 rol-6(su1006gf) was generated by microinjecting 1 ng/µl of WRM0622A_D10 fosmid (https://transgeneome.mpi-cbg.de/transgeneomics/index.html) together with 50 ng/µl pRF4 *rol-6(su1006gf)* into N2. The *xchEx001* strain was then stably integrated into the genome via UV light irradiation and outcrossed 8 times.

We obtained the strain LSD2117 *xchIs016* [P*col-19*::GFP] by injecting 50 ng/µl pJA1 [P*col-19*::GFP] (gift by Ann Rougvie) into N2, followed by UV irradiation and 8 times outcrossing. We UV-integrated collagen overexpressing strains: LSD2013 *xchIs005* [pRF4 *rol-6(su1006gf)* (100 ng/µL)] (=co-injection marker control), LSD2014 *xchIs006* [P*col-13*::COL-13genomic (50 ng/µL); pRF4 *rol-6(su1006gf)* (100 ng/µL)], LSD2017 *xchIs009* [P*col-120*::COL-120genomic (50 ng/µL); pRF4 *rol-6(su1006gf)* (100 ng/µL)], LSD2018 *xchIs010* [P*col-10*::COL-10genomic (50 ng/µL); pRF4 *rol-6(su1006gf)* (100 ng/µL)], LSD1061 was generated by injecting 50 ng/µl pXCH4 (P*col-120*::*col- 120*::dendra2) together with 50 ng/µl pBX *(pha-1 (+))* into LSD9 (*pha-1(e2123*)III). For selection and maintenance of transgenic animals, *the C. elegans* were placed at 25°C.

### Imaging of Matrisome and Adhesome

The genes comprising the *C. elegans* matrisome and Adhesome are curated in Supplementary Table 1.

Unless indicated, animals were kept on NGM plates. For the images, the developmental stages from egg to larval L4 were selected from a mixed plate under the stereoscope and immediately imaged. In preparation for the imaging of day 1 and day 8 of adulthood animals, L4 animals were transferred from NGM plates on plates containing 50 µM FUdR and imaged when they reached their respective age. Depending on whether the fluorescent protein (FP) tag hindered proper secretion and incorporation of the core- matrisome protein, some portion of the FP tagged ECM protein became stuck in the endoplasmic reticulum (ER). In these cases, we largely ignored cytosolic/ER FP signals and focused on FP surrounding plasma membranes that are incorporated into ECM. The fluorescence of the animals was graded on a scale from 0 to 3 intensity. Intensity 3 indicates the highest fluorescence observed. Relative to the highest observed fluorescence of a given reporter line, a gradient scale in 0.5 intervals was categorized and scored, with 0 indicating no fluorescence above the background.

For imaging, we used the BX-51-F Tritech^TM^ Research bright-field fluorescence microscope with a DFK 23UX236 camera, IC Capture 2.4 software, and a triple-band filter from Chroma Technology Corp (described in (Teuscher and Ewald, 2018)). We used 2 mM Levamisole hydrochloride dissolved in the M9 buffer to immobilize the animals for imaging.

### Analysis of collagen-tagged GFP fluorescence intensity

For the analysis of our collagen::GFP strains, we used a Python script written by Elisabeth Jongsma and Jeliazko Jeliazkov in ImageJ (Statzer et al., 2021b). The code is designed to measure the GFP intensity in *C. elegans* animals while ignoring the autofluorescence of the gut. The program takes the area of interest selected from the digital image and compares the intensities for the green and red channels within each pixel. *C. elegans* autofluorescence appears as yellow in the images, a blend of red and green (Teuscher and Ewald, 2018). To remove the autofluorescence without affecting the GFP signal, the red channel intensities are subtracted from the green. Furthermore, signals below a certain intensity threshold are regarded as background noise and also ignored. The program then counts all remaining pixels with intensities in the green channel and adds up the total intensity (it also gives the number of pixels and the mean intensity per pixel). The resulting image can be printed to check if the thresholds were placed properly. The data was further analyzed and visualized using GraphPad Prism 8.2.0. *P*-values were calculated using a two-way ANOVA.

### *In-silico* expression and proteomics analysis

Published datasets were obtained directly from the corresponding supplementary material or through the sequence read archive (SRA). RNA-sequencing datasets were subjected to quality control, quantification (Patro et al., 2017), and subsequent linear modeling (Ritchie et al., 2015; Robinson et al., 2010). Data cleaning and analysis were performed in R (dplyr, ggplot2, clusterProfiler) and using the *C. elegans* Matrisome Annotator (http://ce-matrisome-annotator.permalink.cc/) (Teuscher et al., 2019a).

### COL-19::FRET imaging

*C. elegans* were anesthetized with 25 mM sodium azide (Sigma, S2002-100G) in M9 for live imaging. They were imaged on a coverslip (Menzel Gläser, 24x60mm) covered with a smaller coverslip, which was stuck together with either nail polish (2D or compressed) or with double-sided tape (Sury AG, 3M/9473M25) (3D flow chamber for aging experiments).

*For the formaldehyde crosslinking experiment:* LSD1001 on day 1 of adulthood was fixed in 4% formaldehyde (Sigma-Aldrich, 158127) and imaged the next day. 4% formaldehyde was dissolved in PBS by heating up to 60°C for approximately 2 hours. Formaldehyde solution was sterile filtered and stored at -20°C. We imaged the cuticle as a planar structure in 2D. Therefore 16 μl sodium azide solution was added to a coverslip, and *C. elegans* were subsequently added to this drop. Next, the drop was covered with a smaller coverslip, containing nail polish on each corner for attachment. *C. elegans* prepared with nail polish were imaged at 2048 x 2048 pixel resolution, laser power 20%, zoom 1.5, 400Hz, line accumulation 3.

*For the aging experiments:* We prepared a flow chamber to image *C. elegans* without exerting external mechanical forces (flow chamber, 3D). Therefore 16 μl sodium azide solution was added to a coverslip, and C. elegans were subsequently added to this drop. Next, the drop was covered with a smaller coverslip, containing stripes of double- sided sticky tape (50 µm thickness (3M-VHB; Sury AG, S1473-M25)) on two sides for attachment. *C. elegans* prepared in a flow chamber were imaged at 1024 x 1024 pixel resolution, laser power 25%, zoom 3, 700Hz, line accumulation 4, and z-step size 0.13 μm.

*For chemical manipulation of the FRET ratios during aging:* LSD2052 *pha- 1(e2123)*; COL-19(FRET between exon)-version C3; *spe-9(hc88)* animals were bleached and eggs were distributed on NGM plates to grow at 25°C. At day 4 of adulthood, they were transferred on NGM plates containing 2% ribose (Sigma, R7500- 100G) and nystatin (2.5 ml/1l NGM) (ThermoFisher, 11548886), or 1 mM genipin (Sigma, G4796-25MG), 10 mM MGO (Sigma, 67028-100ML) or 100 mM aminoguanidine (Sigma, 396494-25G). Also, some *C. elegans* were transferred on NGM plates as a control group. After day 4 of adulthood, all *C. elegans* were grown at 20°C and were imaged on day 7 of adulthood. For another approach, LSD2052 animals were placed on day 1 of adulthood on NGM plates containing ribose and nystatin or 1 mM genipin at 25°C. These samples were put at 20°C on day 4 of adulthood and imaged on day 7 of adulthood too.

Imaging was performed with a Leica TCS SP5 confocal microscope (Leica, Microsystems, Mannheim Germany). Cerulean was excited with the 458 nm laser, and emission was detected in the range 470-515 nm (donor channel) and 520-600 nm (acceptor channel). Images were taken with a 63 x PL APO CS 1.4 oil objective (pinhole diameter of 95.5 µm). A maximum intensity projection was performed on z-stacks with ImageJ before quantitative analysis. FRET ratio images were constructed in ImageJ and further visualized and analyzed with LAS AF Lite (Leica Microsystems) and Inkscape. The ImageJ macro for FRET ratio calculation is provided in the last tab of Supplementary Table 12 (FRETanalysisMACRO).

### Imaging and photoconversion of COL-120::Dendra2 and EMB-9::Dendra2 in *C. elegans*

For the RNAi experiments, L4 animals were placed on respective RNAi plates for one generation. To age synchronize the animals, only L4 C*. elegans* of the F1 generation were selected and moved to fresh RNAi plates containing 50 µM FUdR. The animals were imaged on day 2 and day 4 of adulthood, with the photoconversion being performed only on the second day of adulthood. For imaging, they were transferred into a drop of M9 onto 2mm thick, 3% agar pads on microscope slides. For the confocal images, an Olympus FluoView 3000 microscope was used. On day 2, images of the region behind the pharynx and of the vulva region were taken, both before and after the dendra2 photoconversion. Before photoconversion, dendra2’s excitation maxima are at 490 nm and the emission maxima at 507 nm, similar to EGFP; after photoconversion, they change to 553 nm and 573 nm, in the red spectrum. In this study, Dendra2 was photoconverted by using 2% power of the 405 nm laser for 6 sec with 8 µ/s at a resolution of 1024 x 1024. For the analysis, in each image, the *C. elegans* body inside the photoconverted region was manually delineated as a mask, within which the ratio of the total red to green signal intensity is calculated as an indicator of relative amounts of respective proteins. This value was normalized for each image, *i.e.,* per animal and time point, by taking its difference from the similar ratio computed for masks on either side outside the photoconverted region. The data was further analyzed and visualized using GraphPad Prism 9.1.1. *P*- values were calculated using the One-way ANOVA. For details, see Supplementary Table 11.

### C. elegans proteomics

Approximately 1000 - 3000 *C. elegans* were harvested and washed 4 times by centrifugation and resuspension in physiological M9 buffer. Samples were then frozen at -80°C until extraction for all samples in parallel. 500 μL of extraction solution (8 M Urea, 25 mM NH4HCO3, Protease inhibitor (cOmplete tab, Roche Switzerland, 0.25 tablets per 1 ml of extraction solution) and 2 mM Na3VO4, 1mM PMSF) was added to each *C. elegans* pellet on ice. The samples were then processed by bead bouncing for three times 150 seconds using pre-chilled sample holders (-20°C). After centrifugation at 15000 g for 15 minutes (4°C), the supernatant was subjected to total protein quantification and the sample was adjusted to 100 μg of protein. TCEP (Sigma-Aldrich) was added to 5 mM final concentration and the samples were incubated at RT for 30 minutes. Reduced cysteines were alkylated with 10 mM iodoacetamide (Sigma-Aldrich) for 1 hr in the dark at room temperature. Samples were diluted 8x in 50 mM ammonium bicarbonate to reduce the urea concentration to 1 M and protein digestion was performed overnight at 37 C by addition of 2 µg of trypsin (Promega) per sample.

The day after the samples were acidified with the addition of 5% TFA to achieve pH < 3. Desalting was performed using C18 spin columns (Nest group) as suggested by the manufacturer. Columns were wetted with 200 ul (1 CV) of 100% ACN and then equilibrated with 2 CV of 0.1% FA. Following sample loading, the resin was washed three times with 1 CV of 0.1% FA and 5% ACN. Peptides were eluted twice with 0.5 CV of 50% ACN in 0.1% FA and dried under vacuum. The dried peptides of MS-buffer (0.1% FA) with 1:30 iRT peptides (Biognosys) spiked in. The samples were then injected on a TripleTOF 5600 (Sciex, Concord, Canada). Peptides were separated at nano-flow liquid chromatography (NanoLC Ultra 2D, Eksigent) with a flow rate of 300 nL/min using a NanoSpray III source with a heated interface (Sciex, Concord, Canada). The source voltage was 2 kV. The used emitter (Peek 30 cm) was manually packed with 3 μm Reprosil pur (Maisch) beads. The peptides were separated using a 90 min linear gradient from 5% to 30% Buffer B (98% ACN and 0.1% formic acid in HPLC grade H2O) in Buffer A (2% ACN and 0.1% formic acid in H2O).

For data acquisition, the instrument was operated in positive ion, with high sensitivity SWATH-mode using 64 variable-width windows precursor isolation scheme, between 350 and 1500 m/z with a 1 m/z one-sided overlap. The Updated SWATH-window scheme is essentially described by Collins et al. 2017 (Collins et al., 2017). Accumulation time was set to 250 ms for the precursor survey scan and 50 ms for each of the 64 MS2 fragment ion scans, which resulted in a total cycle duty time of 3.5 s. Dynamic collision energy (CE) and collision energy spread (CES) was optimized for fragmentation of each peptide following the rolling collision energy formula (CE= m/z * Slope + Intercept) with a collision energy spread of 15 eV. The SWATH data was searched in Spectronaut v13 (Biognosys) using a *C. elegans* FASTA (4352 entries) and directDIA with default BGS settings. Downstream analysis was performed in R (Zhang et al., 2018). The raw files and search results are available in PRIDE under the identified PXDXXXXXXXX

### RNAi clones and libraries

#### Generation of RNAi clones

We cloned the 942 bp *col-120* cDNA into pL4440, validated correct insertion and sequence, and transformed this plasmid (pLSD051) into HT115.

#### Generation of RNAi screening libraries

For the target RNAi screen, we worked with 5 RNAi bacteria libraries containing selected RNAi clones to knock down specific gene classes or categories. The kinase library, two transcription factor libraries (bZip and TXN- factor Libraries), and the metabolism library were a generous gift from Gary Ruvkun (Harvard Medical School) (Venz et al., 2020). We constructed our Matrisome library based on our definition of the *C. elegans* matrisome (Teuscher et al., 2019a). The library contains 652 RNAi clones of the 719 *C. elegans* matrisome genes. For the missing 67 genes, no RNAi clones were available. The bacteria were picked from either the ORF-RNAi or the Ahringer RNAi libraries (both available from Source BioScience). Bacteria glycerol stocks of the clones were transferred with a pipette into 96-well plates each also containing control wells with control bacteria RNAi clones (L4440 (empty vector control), *daf-2*, *bli-3*, *daf-16*, *skn-1*, *gfp*, *col-144*) and some empty wells (LB mixed with glycerol).

The same procedure was followed for generating the two validation screen libraries. Validation library I consisted of hits from the previous screen rounds, together with selected clones of hits from the P*col-12*::dsRed expression screen performed by the Ewbank lab (Zugasti et al., 2016) and selected clones of genes from our literature research. Validation library II consisted of hits from the Validation library I screen.

### RNAi screen

The screen was performed on 96-well plates, each well containing 150 μl NGM with 100 μg/ml Ampicillin and 1mM IPTG, seeded with 8 μl concentrated RNAi bacteria. The bacteria were grown overnight in 96-deep-well plates in 800 μl LB containing 100 μg/ml Ampicillin and 12.5 μg/ml Tetracycline. The next morning, another 700 μl LB with Ampicillin and Tetracycline was added. After four additional hours of growth, the plates were centrifuged, the supernatant was discarded, and each well was filled up with 35 μl LB containing 100 μg/ml Ampicillin and 1mM IPTG to seed the 96-well NGM plates. For the preparation of the *C. elegans*, we used plates containing gravid adult *C. elegans*. We age synchronized the animals at stage L1, by dissolving the parent but leaving the eggs intact and letting them hatch in an M9 medium with cholesterol, without food, so they stage arrest until all eggs were hatched (Teuscher et al., 2019b). The next day, we placed 25 animals of our screening strains in wells of 96-well containing Normal Growth Medium (NGM), each plate was seeded with one of the clones from the 96-well library plates. *C. elegans* grew up at 25°C until day 2 of adulthood, as this temperature is needed to activate *glp-1* and *spe-9* mutations during development and make them sterile, later moving them to 20°C. The plates were scored on adulthood day 1 and day 8 under a fluorescence stereoscope and the fluorescence of the animals was graded on a scale from 0-3 intensity. The result was counted as a hit when the average of the three to four replicates was at least 0.5 (lowest visible difference) over the control. For the gene ontology enrichment, WormCat was used (Holdorf et al., 2019).

### Time-course measurements of fluorescent expression reporter strains

The strains for time-course measurements were placed as L4s on plates containing 50 µM FUdR and were scored on the indicated days under a fluorescence stereoscope. For each animal, the fluorescence was graded on a scale from 0-3 in 0.5 steps. Per measuring round 20-30 *C. elegans* were used. The data was further analyzed and visualized using GraphPad Prism 8.2.0. P-values were calculated using a two-way ANOVA.

### EMB-9/PAT-3 co-localization experiments

For the imaging of a potential co-localization of the EMB-9::mCherry and PAT-3::GFP in *C.elegans*, the animals were placed as L4s on plates containing 50 µM FUdR. Images were taken on day 1, day 3, and day 8 of adulthood using an Olympus FluoView 3000 microscope. For the RNAi experiments, the animals were placed as eggs on plates seeded with control L4440 RNAi or *daf-2* RNAi bacteria. The RNAi NGM plates contained 100 μg/ml of Ampicillin and 1mM IPTG. The L4 animals were moved to RNAi NGM plates that additionally contained 50 µM FUdR.

For the analysis of colocation, we separately assessed this for the body wall muscles around the head region, midbody region, and tail region, by imaging and manually selecting these regions for each *C. elegans* (all images and area selections are provided in Supplementary Table 11). In our study, *colocation* is defined as very close proximity (beyond the pixel resolution of the employed microscopy imaging) of the proteins and hence their fluorescence marker signals. To that end, red and green intensity (signal) at each pixel indicates the amount of the corresponding protein within the space covered by that image pixel. If these intensities are *similar* for the same pixel, this would mean similar amounts of each molecule within that pixel (thus in “very close proximity” per definition). Instead of the concept of similarity, i.e. being “equal”, we chose to quantify their correlation (*i.e.,* the null-hypothesis of them being in relation with a fixed ratio) because intensity equality between red and green markers would require strict constraints, such as the imaging system scaling raw reading similarly to all RGB values, these different markers reacting the exact same way to optic excitation, each marker staining the molecules in the same fashion, etc. Instead, a correlation analysis checks for an arbitrary (linear) relation model between these markers occurring together in pixels (i.e., colocating). Accordingly, we computed the correlation of per-pixel red-to-green signal across the image, per animal, and time point. All pixel quantifications and calculations are provided in Supplementary Table 11. Since the structures we aim to quantify are smaller than the pixel size, the intensity red/green quantifies how many/much collagen and integrin exist in each pixel. Then, observing more green where the red is, and vice versa (*i.e.,* the correlation metric we used) indicate their co-occurrence spatially. The term “disassociation” refers to our quantification of the loss of spatial relationships between the collagen (red) and integrin (green).

### Lifespan assays

Manually lifespan assays were performed as described in (Ewald et al., 2016). In brief, L4 animals were picked onto culturing plates containing 50 µM FUdR. TJ1060 *spe- 9(hc88)* I; *rrf-3(b26)* II and CB4037 *glp-1(e2141)* III were grown at 25°C until day 2 of adulthood and then placed on RNAi plates (without FUdR) at 20°C for the remainder of the lifespan. The *spe-9(hc88)* is a temperature-sensitive sterile mutation (L’Hernault et al., 1988). SPE-9 is a transmembrane protein on the sperm required for oocyte interaction (L’Hernault et al., 1988; Zannoni et al., 2003).

Lifespan machine assays were performed as described in (Statzer et al., 2020; Williams et al., 2021). In brief, L4s were washed onto RNAi culturing plates containing 50 µM FUdR until day 5 of adulthood at 20°C, and then placed on special RNAi plates and placed into the lifespan machine. For experimental details, setup, raw data, and statistics, see Supplementary Table 8.

Comparative automated lifespan analysis: To compare the longevity-promoting effects of *daf-2* RNAi on different *C. elegans* matrisome mutants, the mean lifespan of *daf-2* RNAi was plotted against the corresponding mean lifespan on L440. N2 data from multiple lifespan assays was compiled, representing the general lifespan extension of *daf- 2* RNAi in N2. For the analysis, the different strains were grouped according to their respective matrisome category. The code can be found in Supplementary Table 7 and on Github (https://github.com/Ewaldlab-LSD/DiagonalPlots).

*For compound treated lifespans:* TJ1060 *C. elegans* were bleached and the eggs were distributed on NGM plates to grow at 25°C. On day 4 of adulthood, 100 *C. elegans* were placed on 10 NGM plates containing 2% ribose and nystatin and another 100 on 10 control plates. LSD2052 worms were also bleached, and eggs were distributed on NGM plates at 25°C. This time, 50 *C. elegans* were transferred on two plates with 1 mM genipin, 50 on two DMSO (90.4 μl DMSO/10ml NGM) (Aldrich, M81802) plates and another 50 on two control plates. All samples were put at 20°C on day 4 of adulthood. In the following days, plates were checked for dead animals every day. Lifespan experiments with aminoguanidine were performed by using the lifespan machine. N2 worms were bleached and incubated in M9 for 2 days at 20°C on a rotor to synchronize the animals at the L1 stage. L1 animals were then transferred on NGM plates and grown to larval stage L4. From that time on, nematodes were grown on NGM plates containing FUdR. On day 4 of adulthood, we transferred approximately 30 worms on one plate, 4 plates in total per experimental condition. The plates were prepared with 1 mM, 50 mM, and 100 mM aminoguanidine and control plates without any interventions. All plates contained FUdR and nystatin.

### Quantification of cuticle thickness using electron microscopy images’

Transmission electron microscopy (TEM) images of *C. elegans* age day 2, day 7, and day 15 were downloaded from wormimage.org. Only transverse sections were considered as they were thought to minimize the error in measurement due to oblique cutting angles. Regions distorted by the cutting process were excluded from the analysis. Suitable regions were manually traced and the resulting binary masks (colored curves, one pixel thick) were exported to Matlab (Supplementary Figure 5 A-C). As the borders of every cuticular layer were traced separately, both the total thickness and thickness of the individual layers could be measured.

To determine the mean distance between two-layer borders (i. e. the mean thickness of the cuticle layer) represented by two binary masks A and B, the euclidean distance transform of mask A was calculated leading to Admap. The Euclidean distance transform of A assigns to every pixel in A the distance to the nearest non-zero pixel. By element-wise matrix multiplication of Admap and B, the nearest distance to A of every pixel in B can be calculated leading to vector b (Supplementary Figure 5 D). By repeating the process while swapping A and B, the closest distances to B for every pixel in A can be calculated. The distance between two curves A and B was then defined as the mean value of the shortest distances to the other curve for every pixel in both A and B.The dataset used contained digitized TEM images of various different magnifications (*m*) (1200x – 15500x). Every image contained a scale bar of the length of 2.5 cm. For analysis, the length of the scale bar in pixel (*s* [px]) was measured. The conversion factor (*f* [μm/px]) was calculated according to formula (1) and multiplied with the final measurements to convert them to micrometers.

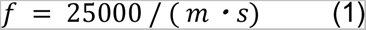

### Fluorescence microscopy of YAP-1::GFP expression

Strain YAP-1::GFP was maintained on OP50 NGM plates at 20⁰C. Gravid adults were treated with sodium hypochlorite solution to obtain eggs. The eggs were grown on L4440 and *daf-2* RNAi plates until the L4 stage. At L4, the plates were transferred to 25⁰C or kept at 20°C. Then, at day 1 of adulthood, approx. 30 animals were mounted onto a 2% agarose pad and anesthetized with 20 mM levamisole for imaging. Animals were captured at 10X with one or two fields of view and then, stitched together later, using ImageJ. Quantification of the total fluorescence intensity was done by running a python script, GreenIntensityCalculator (available publicly in Github-Ewaldlab: https://github.com/Ewaldlab-LSD) in ImageJ (Statzer et al., 2021a). Statistical analysis of the quantified data was done using graph pad prism software (GraphPad Prism 9.0). The experiment was performed in three independent biological batches.

### Fluorescence microscopy for YAP::GFP expression on hemidesmosome- containing structures

Strain *ihIs35* YAP-1::GFP was maintained on OP50 NGM plates at 20⁰C. Gravid adults were treated with sodium hypochlorite solution to obtain eggs. The eggs were grown on L4440 and *daf-2* RNAi plate until young adults. At the young adult stage, the plates were transferred to 25°C. Then, on day 3 of adulthood, approx. 30 animals were mounted onto a 2% agarose pad and anesthetized with 20 mM levamisole for imaging. Animals were captured at 100X with one posterior field of view. Quantification of the distribution of the categories of hemidesmosome-like phenotype was done manually. Category 0-1 were allotted to absent to faint appearance of hemidesmosome-containing structures (as in most L4440 control conditions) while 2-5 category represent the graded intensity of the present hemidesmosome-containing structure (as observed to be more the case in *daf-2* RNAi condition). Statistical analysis of the quantified data was done using graph pad prism software (GraphPad Prism 9.0). The experiment was performed in four independent biological batches.

### Confocal microscopy for colocalization of YAP-1::GFP with VAB-10::mCherry

Transgenic strain LSD2207 YAP-1::GFP;VAB-10A::mCherry was grown on L4440 or *daf- 2* RNAi plates at 20°C. At young adult stage, animals were shifted to 25°C. After 24h, animals were mounted onto 2% agarose pad and anesthetized with 20 mM levamisole for confocal imaging. Animals were captured with confocal laser scanning microscope (Olympus Fluoview FV3000), using 60X oil objective. Images were taken at an intensity of 0.7% for red channel, and 30% for green channel, at 4.00 zoom, at resolution of 1024 × 1024 pixels and with a gain of 1.0. The experiment was performed in three independent biological trials.

### Pressure application and quantifying GFP intensity

Approximate 30 L4 animals were picked onto the center of a 60 mm OP50 FUdR (no FUdR for LSD2002 animals because containing temperature-sensitive sterile mutation *spe-9(hc88)*) containing plate, for each condition in technical replicates of two. Then, with the help of a sterilized spatula, a 2 cm x 2 cm chunk of agar was cut and transferred to a glass slide gently. After that, another glass slide was pressed upon this agar chunk gently, sealing the glass slides with the help of tape. For the pressure condition, an inverted 50 mL water-filled falcon was placed on top. The approximate pressure was calculated as the force divided by the area. Thus, the pressure was (0.05 kg weight x 9.81 m/s^2^ (=g))/ (0.04 m^2^ agar area) = 12.3 Pa. This pressure device was kept at 20°C or 25°C as indicated in the figure legend for three days, after which the animals were imaged. To assess fluorescence intensity, the agar chunk was transferred from the pressure device to a fresh OP50 plate and let the animals move out of the agar for about half an hour. Animals were then mounted onto a 2% agarose pad and anesthetized with 20 mM levamisole for imaging. They were captured at 10X with one or two fields of view and then, stitched together later, using ImageJ. Quantification of the total fluorescence intensity was done by running a python script, GreenIntensityCalculator (available publicly in Github- Ewaldlab: https://github.com/Ewaldlab-LSD) in ImageJ (Statzer et al., 2021a). Statistical analysis of the quantified data was done using graph pad prism software (GraphPad Prism 9.0).

